# Single-and double-strand circulating DNA fragmentomics for enhanced cancer detection performance

**DOI:** 10.1101/2025.09.17.676909

**Authors:** Ekaterina Pisareva, Marianne Richaud, Severine Tabone-Eglinger, Thomas Bachelot, Benoît Roch, Thibault Mazard, William Jacot, Veronique Pezzella, Benjamin Besse, Fabrice André, Marc Ychou, Jacques Colinge, Alain R. Thierry

## Abstract

In early detection of cancer, the use of circulating cell-free DNA (cirDNA) obtained from blood samples is notable for its minimally invasive nature. We have developed an algorithm designed to discriminate cancer patients and healthy individuals based on cirDNA fragment end motif analysis assisted by machine learning, using data obtained from shallow whole genome sequencing (a method we call EMA). We applied EMA to cirDNA from the plasma of patients with stage II-III breast cancer, stage I-III non-small cell lung cancer, and metastatic colorectal cancer (mCRC). CirDNA from 158 individuals was prepared following the conventional double-stranded DNA library preparation (DSP). Using 3 bp end motifs, each tumor type was detected with a sensitivity of 0.87-1.00, a specificity of 0.95, and an AUC above 0.96. The three selected cancer types could be differentiated with an accuracy (ACC) above 0.94. Multi-cancer detection by pooling samples from the three cancer types showed ACC, AUC and sensitivity of 0.98, 0.99 and 0.98, respectively. Comparisons with 4 and 2 bp end motifs were conducted, and our main observations were confirmed using an external public dataset (N=366). We also performed a single-stranded DNA library preparation (SSP) using mCRC patients and healthy control cirDNA, which allowed us to make the first ever end motif analysis in the literature which compares the use of DSP and SSP. As compared to EMAD (use of DSP), EMAS (use of SSP) produced a very significant difference in end motif frequency and an improved cancer detection performance (ACC, AUC and sensitivity of 0.97, 1.00 and 0.99, respectively). Furthermore, optimal performance was produced when the full-size range was used for EMAS, whereas when the dataset was restricted to fragments of 115 - 220 bp for EMAD. EMAS strong performance, coupled with its compatibility with cost-effective shallow cirDNA sequencing, positions this methodology as a potentially transformative tool in early cancer screening.

## INTRODUCTION

Efforts to improve early cancer detection have typically focused on a range of approaches, both invasive and non-invasive. Within the latter category, blood plasma has gained significant attention as a readily accessible biological medium, which can provide real-time insights into tumor biology ^1,2^. However, the lack of well-established biomarkers by which specific cancers can be identified has proved problematic, especially with early-stage cancers ^3^. In this context, circulating cell-free DNA (cirDNA) represents a novel source of information that offers unique advantages for non-invasive diagnostics ^4–8^. Recently published work has shown that the study of cirDNA fragments (called fragmentomics), while still in its infancy, has the potential to become a powerful diagnostic tool. Indeed, we and others have already demonstrated the link between cirDNA fragments and disease states, providing robust proof-of-concept data for this innovative approach ^4,9–13^.

CirDNA fragmentation derives from chromatin packaging around mononucleosomes. Nucleosomes protect DNA against nucleases present in cell cytoplasm upon DNA release and in the circulation subsequently ^4,14^. Arising from these mechanisms, cirDNA fragments features are the product of nuclease digestion and of the protection provided to DNA as a result of its interactions with mononucleosomes. Therefore, cirDNA fragments observed in blood plasma constitute the net product of nuclease digestion and the protection provided to DNA via its physical interactions with mononucleosomes. The field of fragmentomics has investigated different properties of cirDNA towards cancer detection or for a deeper understanding of its fundamental mechanisms. Among the first features of cirDNA to be exploited were the presence of cancer-associated mutations from DNA released from tumor cells, thereby enabling tumor detection ^2,3,15^ and the follow-up of residual disease post-therapy ^16^. Another standard area of investigation in this field is the association of particular cirDNA fragment lengths distribution with the presence of cancer, which we and others have pioneered ^17–19^. More recently, additional characteristics of cirDNA have gained growing attention. Amongst others, these include cirDNA methylation status ^20–22^, the analysis of DNA end motifs found at the extremities of cirDNA fragments ^23,24^, and the occupancy of transcription factor binding sites ^25^. The emergence of inexpensive DNA sequencing technologies, along with the application of specific machine learning (ML) and artificial intelligence (AI) methods, has considerably accelerated the development of fragmentomics ^26,27^. Notably, it has enabled numerous studies to link fragmentomics properties with cirDNA fragment genome mapping.

Here, we present a new perspective on cirDNA end motif analysis associated with cost-effective shallow whole genome sequencing (sWGS), specifically in relation to the detection of early stage cancer. By applying ML to an analysis of the relative frequencies of end motifs in each sample, our EMA method showed highly sensitive and specific cancer detection capabilities. In parallel, we employed both the commonly used double-stranded DNA library preparation (DSP) protocol and the less frequently used single-stranded DNA library preparation (SSP) ^17,28,29^ on HI and mCRC. This allowed us to explore differences in end motifs at 5’ and 3’ cirDNA fragment extremities, following EMA using DSP (EMAD) or SSP (EMAS) and to test whether the cancer detection power of EMAS was comparable to that of EMAD (Fig. 1). In addition, we explored the relationship between natural cirDNA fragment size ranges and the performance of both EMAD and EMAS. EMA is very flexible with respect to the choice of the particular ML model used, and is compatible with DSP and SSP data, even when subjected to specific fragment size ranges.

**Figure 1.**
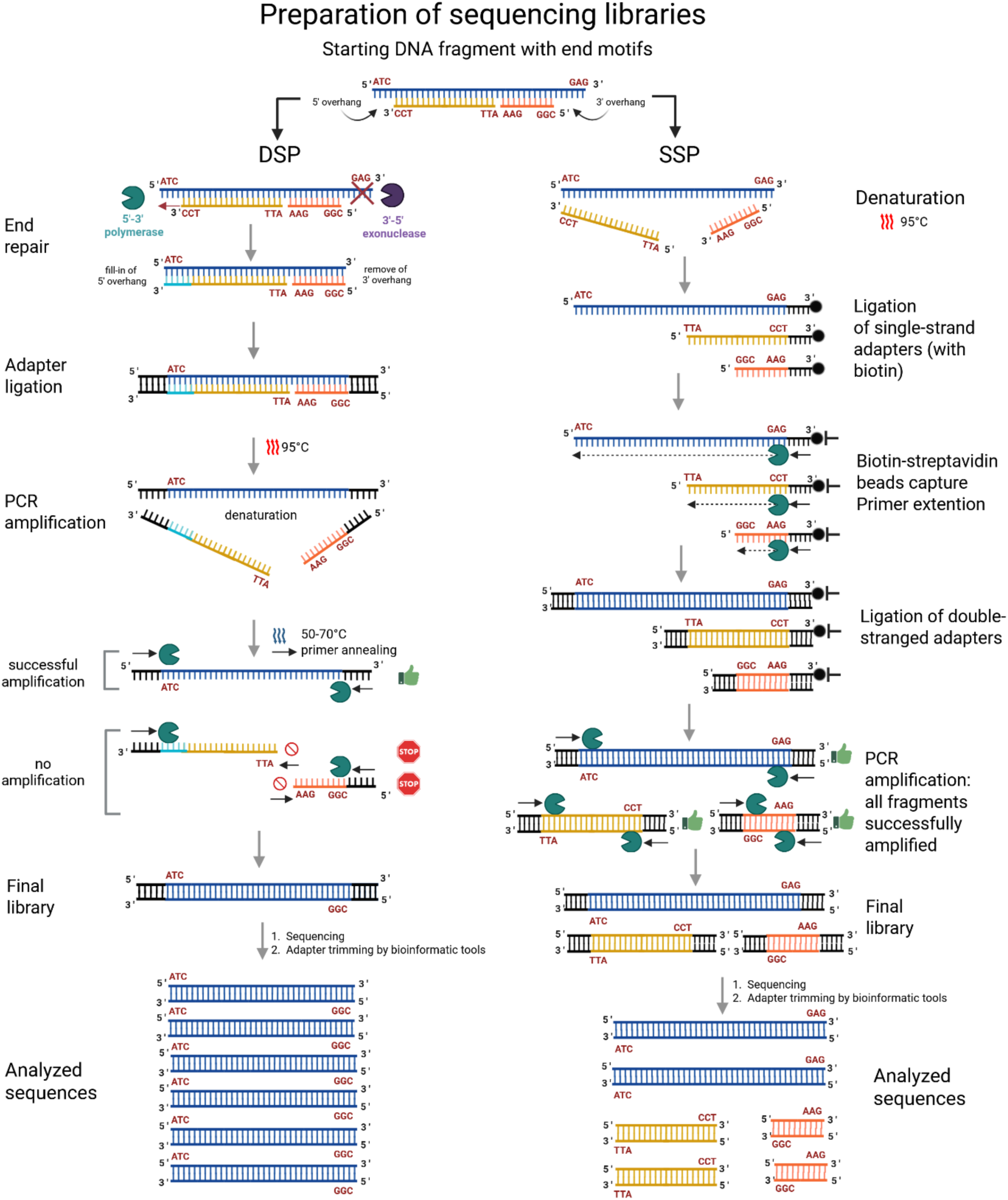
Schematic illustration of key steps in the DSP and SSP library preparation protocols used in this work and their sensitivity to recover end motifs of DNA fragments with overhangs and single-strand breaks.

**Figure 1.**
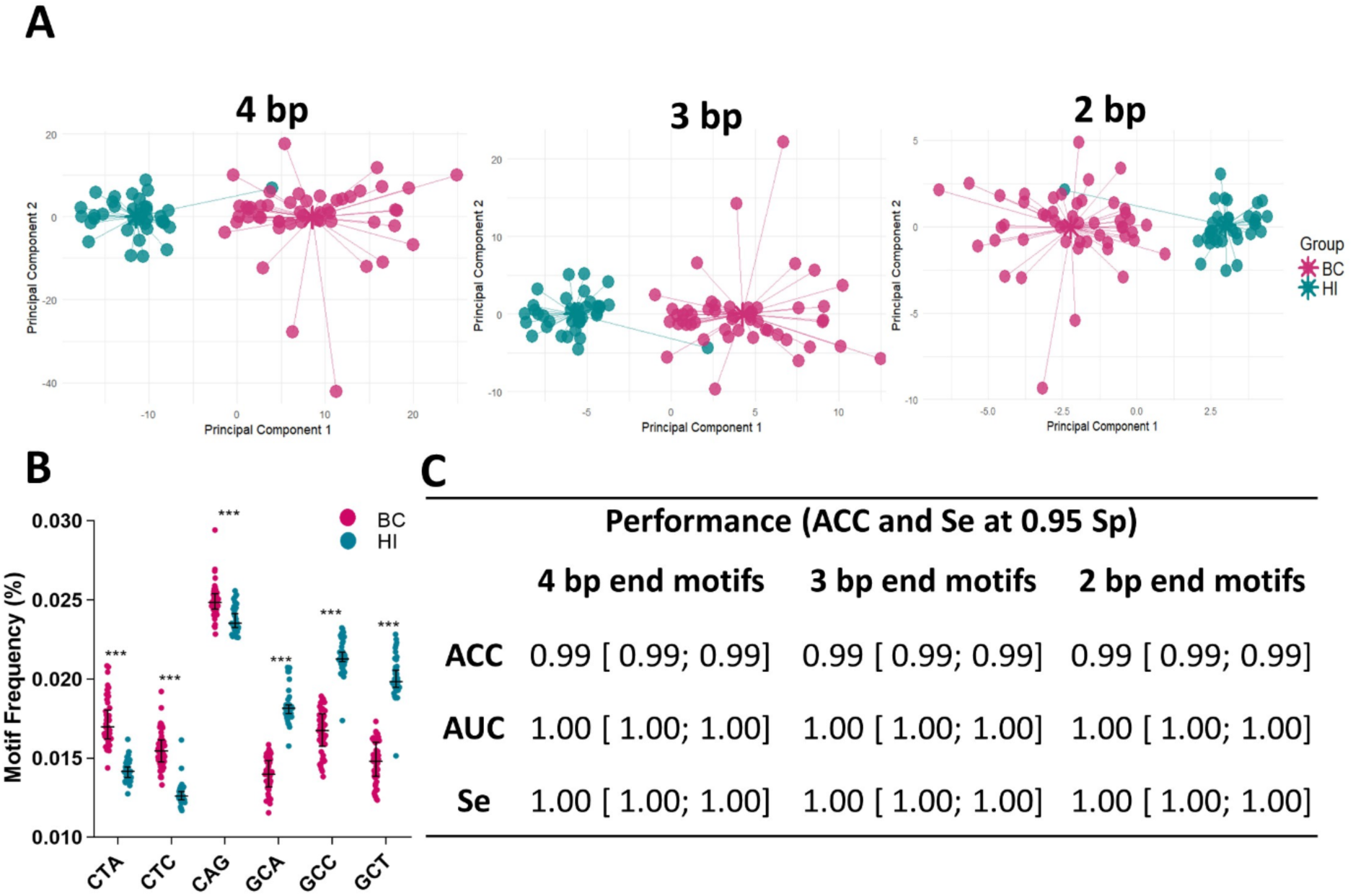
EMAD in breast cancer. **(A)** Two-dimensional PCA projections of 4, 3 and 2 bp end motif relative frequencies in plasma cirDNA fragments of 37 healthy individuals (HI) and 50 breast cancer (BC) patients (stage II–III). **(B)** Six representative 3 bp motifs showing significant relative frequency biases between BC and HI subjects. Wilcoxon 2-sided test with Benjamini-Hochberg correction. **(C)** EMAD performance combined with Random Forest for BC detection (5-fold CV). We imposed specificity at 0.95 to determine sensitivity and accuracy. ACC = accuracy, AUC = area under the receiver operating characteristics curve, Se = sensitivity, Sp = specificity, reported intervals are 95% confidence intervals.

## RESULTS

### Definition of end motifs of cirDNA in plasma

We analyzed the end motifs of cirDNA fragments in plasma using sWGS of isolated cirDNA following DSP-sWGS or SSP-sWGS protocols (Fig. 1). End motifs were identified using the first 4, 3, or 2 nucleotide sequence on each 5ʹ fragment end of plasma DNA after alignment to the human genome. In some cases, we also exploited 3’ fragment end motifs. Unless explicitly specified, only the 5’ end motifs were used. Given that there are four possible nucleotides (A, G, T, and C), the number of end motifs in 5’ or 3’ was 4^𝑛^, with 𝑛 the chosen motif length. The relative frequency of each end motif was computed for each sample by dividing an end motif frequency by the total number of end motifs identified in the sample.

### Early-stage breast cancer detection using cirDNA end motif analysis from DSP-sWGS data

To study whether the relative frequencies of 4, 3, and 2 bp end motifs would differ in patients with cancer, we sequenced the plasma cirDNA of 37 HI and 50 patients with stage II-III breast cancer (BC). Principal Component Analysis (PCA) was used to evaluate whether BC subjects displayed identifiable characteristics of end motifs at each motif length considered. Confirming this suspicion, we did indeed find that BC subjects tended to cluster together, while HI subjects also formed a distinct if more compact cluster independent of motif length (Fig. 2A). Comparison of the cirDNA end motif relative frequencies of the BC and HI groups revealed a number of significant differences. Figure 2B features six representative 3 bp motifs, including three motifs (CTA, CTC, and CAG) for which there was a significant increase in BC patients, and another three motifs (GCA, GCC, and GCT) with a significant decrease in BC patients.

**Figure 2.**
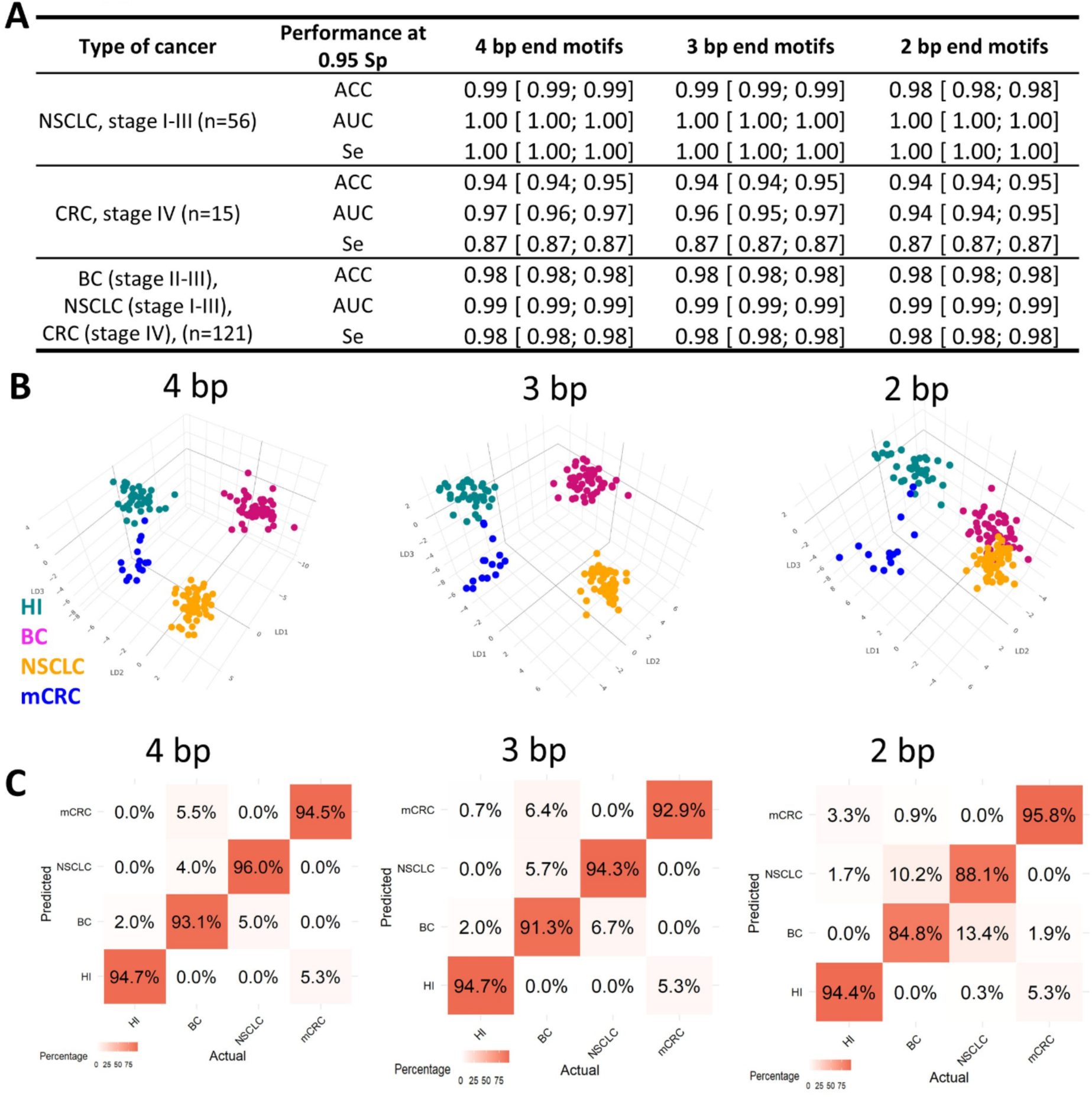
Modulation of end motifs of cirDNA fragments across different cancer types. **(A)** EMAD with Random Forest (RF) models for detection of non-small cell lung cancer (NSCLC), metastatic colorectal cancer (mCRC), and a combined group of all three cancers versus 37 healthy individuals based on 2-4 bp end motifs relative frequencies. Model performance was evaluated by 5-fold cross-validation and imposing 0.95 specificity. **(B)** Three-dimensional projection following linear discriminant analysis (LDA) of 2-4 bp end motif relative frequencies. We note clusters of BC, NSCLC, mCRC, and healthy individual (HI) samples. **(C)** Confusion matrices showing the performance of RF model classifying BC, NSCLC, mCRC, and HI samples based on 2-4 bp end motif relative frequencies. The matrix is the average over the test sets of 5-fold cross-validations repeated 100 times.

To explore whether, with the aim of detecting patients with cancer, an automatic classifier could be designed which might exploit end motif relative frequencies, we trained and tested a Random Forest (RF) model ^30^ for each motif length. It involved determining end motif relative frequency computation from EMAD. Five-fold stratified cross-validation (CV) repeated 100 times enabled us to estimate EMAD accuracy (ACC), specificity (Sp), sensitivity (Se), and area under the receiver-operating characteristic curve (AUC). In cancer detection methods, as a general rule, both to maintain cost-effectiveness and to reduce the psychological burden of screening in large populations, high specificity is preferred. We hence reported RF model ACC and Se imposing a minimum Sp at 0.95. For that model, we obtained Se of 1.00 and ACC of 0.99, regardless of the length of cirDNA end motifs considered (Fig 2C).

### Applicability to lung and metastatic colorectal cancers

To confirm the broad applicability of EMAD in cancer detection, we sequenced plasma cirDNA samples from two additional cohorts. The first cohort consisted of patients with stage I-III non-small cell lung cancer (NSCLC, n = 56), and the second cohort consisted of patients with metastatic colorectal cancer (mCRC, n = 15). As above, EMAD was applied with RF, and its performance estimated using 5-fold CV (Fig. 3A). Whilst stage I-III NSCLC plasma samples displayed ACC, AUC, and Se values equal to 0.99, 1.00, and 1.00, respectively, for 3 bp end motifs, mCRC plasma samples showed slightly lower values: AUC 0.96, ACC 0.94, and Se 0.87. As for BC, no real variation in performance was observed between different motif lengths.

**Figure 3.**
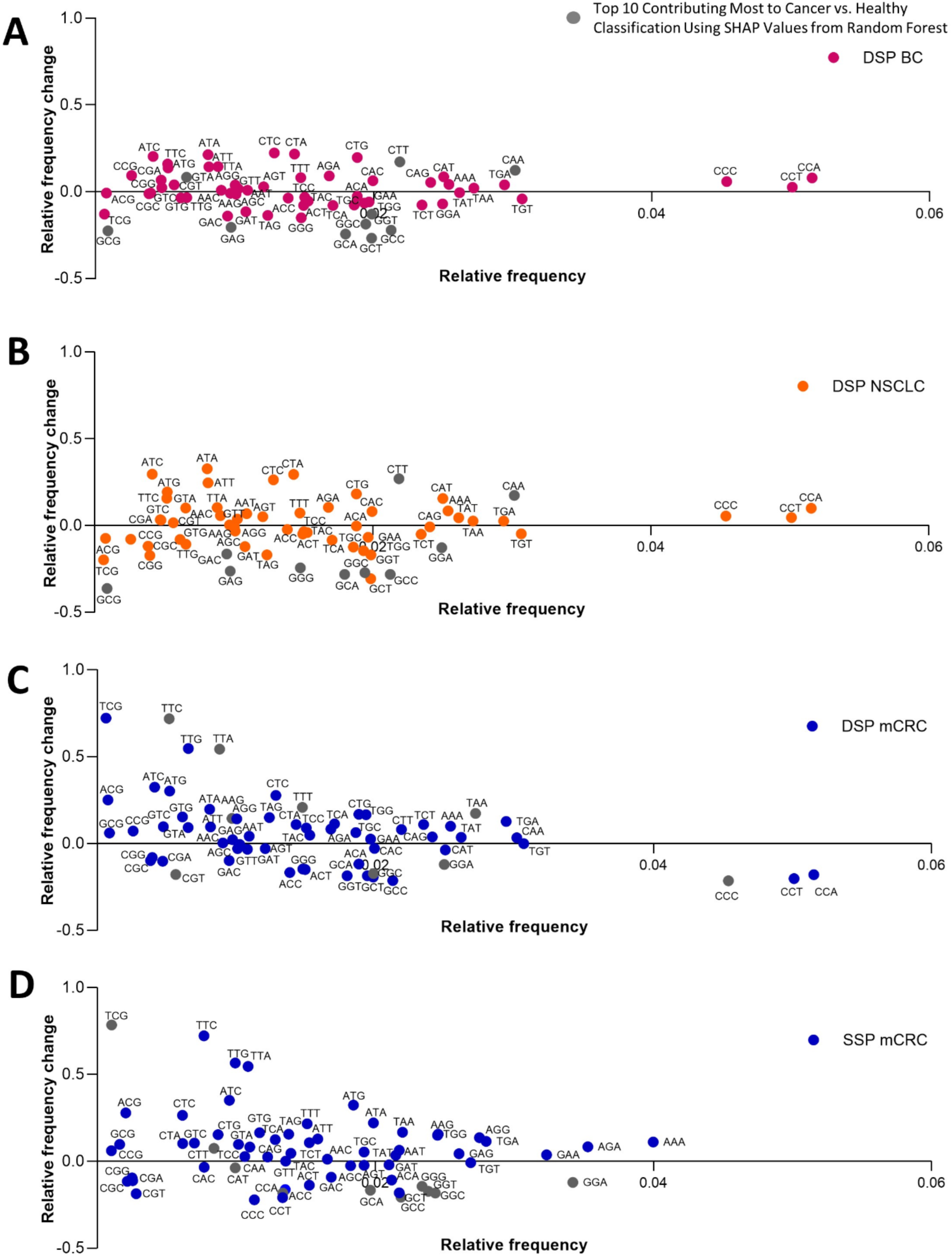
Changes in the relative frequency of 3 bp end motifs in patients with: **(A)** breast cancer (BC), **(B)** non-small cell lung cancer (NSCLC) and **(C)** metastatic colorectal cancer (mCRC) analyzed using DSP and **(D)** mCRC analyzed by SSP as compared to healthy individuals (HI), shown as a function of motif frequency in HI.

Our next endeavor was to assess whether EMAD could distinguish between the three cancers considered in this study. This possibility was clearly supported by the results of linear discriminant analysis (LDA)-based 3-dimensional projections for the different motif lengths (Fig. 3B), albeit with reduced potential where 2 bp end motifs were employed. RF models were trained and tested by 5-fold CV to specifically quantify EMAD’s ability to distinguish the three cancer types. Our results reveal the accuracy of its cancer classification as > 0.93 for 4 bp motifs, > 0.91 for 3 bp, and > 0.85 for 2 bp (Fig. 3C). It was noticeable that, in the evolution from 2 to 3 bp, there was a greater increase in performance than in the evolution from 3 to 4 bp. This would appear to imply that the 3 to 4 bp motif ends may capture more information related to fragmentation signatures.

Another implication of EMAD’s ability to distinguish between different cancer types is that it points to the existence of cancer-specific signatures. To properly evaluate this phenomenon, we first assessed 3’ and 5’ end motif frequency differences. Using DSP, we expected no major differences, given that end repair is an integral part of the DSP protocol, and indeed this proved to be the case, as illustrated in HI (Fig. S1A) and our cancer cohorts (Fig. S1B, Fig.S2). Comparing HI, BC, NSCLC, and mCRC 5’ end motif frequencies we found significant differences in a majority of motifs, especially in mCRC (Fig. 4 and Fig. S1B and S2). In terms of magnitudes, most differences were rather modest (<10%) but RF could nonetheless capture recurrences within a cohort and deliver good discriminatory power (Fig. 3C).

**Figure 4.**
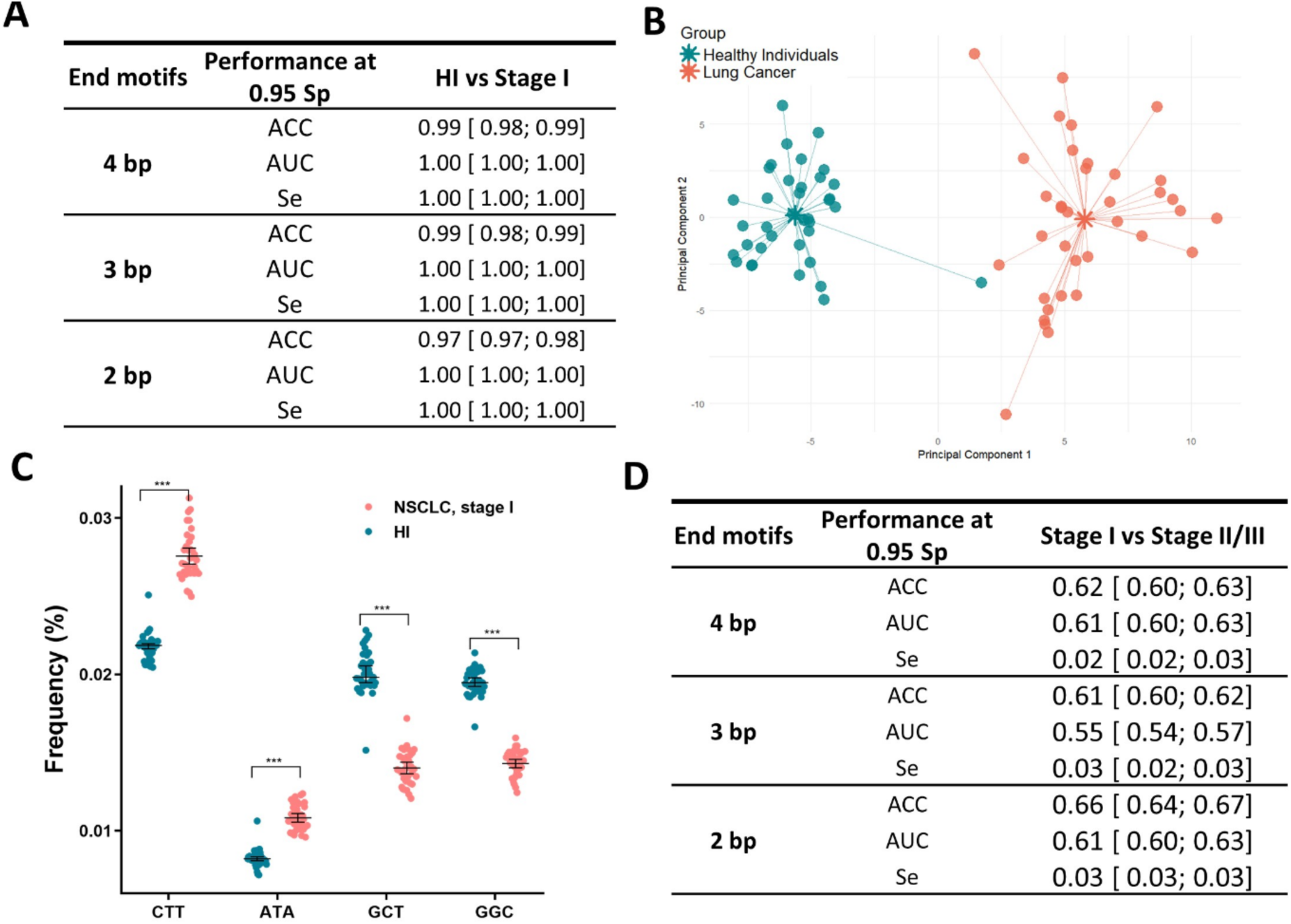
EMAD on stage I non small cell lung cancer. **(A)** EMAD performance with Random Forest (RF) models for classification of stage I of non-small cell lung cancer (NSCLC) and healthy individuals (HI) using 4, 3 and 2 bp end motif relative frequencies. Model performance was evaluated using 5-fold cross-validation and imposing 0.95 Sp. **(B)** Two-dimensional projection by PCA of 3 bp end motif relative frequencies showing almost perfect separation of stage I NSCLC from HI. **(C)** Four representative 3 bp motifs displaying significant differences in relative frequencies between stage I NSCLC and HI. **(D)** Performance of the RF models for classification of stage I versus stage II/III of NSCLC using 4, 3 and 2 bp end motifs.

While EMA considers all the end motifs as a whole providing strong classification through ML, it is worthwhile to individually study the end motif frequency and relative frequency change in comparison to controls to investigate cirDNA fragmentation at the deoxynucleotide level and to compare it to the previously reported other methods. The end motif sequences with the highest relative frequencies following DSP-sWGS for the three studied cancer types were the same: CCC, CCT, and CCA (Fig. 4A-C). Inspection of these three motifs led to positive relative frequency difference in BC and NSCLC samples while leading to negative difference in mCRC samples. The motifs with the lowest relative frequencies following DSP for the three cancers studied were ACG, TCG, and GCG. However, these three motifs showed a positive change in relative frequency for mCRC, unlike breast and lung cancer (Fig. 4B). The end motifs that showed the greatest positive difference in relative frequency were ATC, ATA, CTA, CTC, and CTA for lung and breast cancer; whereas it was TCG, TTC, TTG, TTA and ATC for mCRC. The end motifs that showed the largest negative difference in relative frequency were GCG, GCT, GCA, GCC and GGC for breast and lung cancer (Fig 3A and B), while it was CCC, GCC, CCT, CGT and GCT for mCRC (Fig. 4C). Within EMAD, the use of SHAP values from Random Forest allows the level of contribution of each end motif to discriminate between HI and cancer to be determined (Fig. S5). As illustrated by end motifs showing the top 10 SHAP values (Fig. 4), there is a moderate concordance between relative frequency change as determined following DSP-sWGS and contribution to the classification following EMAD. Whereas end motifs may be different, the end motifs with the highest SHAP values are among those with the highest relative frequency change (Fig. 4).

As determined by SHAP analysis (Fig. S5) from Random Forest classification indicates that some end motifs are specifically efficient in distinguishing a cancer type: GTA and GGA are specific to distinguish BC; GGT,CGG, and ATT to distinguish NSCLC; and TTT, TTG, TTC, TTA, TGA, TCG, TAA, CGT, CCT, CCC, CCA, CAA, ATA, AGC, ACT, ACG, ACC, and AAG to distinguish mCRC.

The top10 more elevated SHAP values using EMAD for BC are GCT, GGC, GCG, GCC, GCA, GAG, GTA et GGT, CTT, CAA (Fig. S5), while CCA,CCT,CCC,CAA, CAT, TGA,TAA and TGT showed the highest level of relative frequency (Fig. 4). The top10 more elevated SHAP values using EMAD for NSCLC are GGG, GGA, GCG, GCC, GCA, GAG, CTT, CAA, GGC and GAC, while CCA,CCT,CCC, CAA, CAT, TGA,TAA and TGT showed the highest level of relative frequency. The top10 more elevated SHAP values using EMAD for mCRC TTT,TTC,TTA,TAA, GGG, GGT, GGA, CCC,CCA, CGT, and AAG while CCA,CCT,CCC, CAA, TGT, TGA,TAA and TAT showed the highest level of relative frequency.

To evaluate EMAD’s detection of stage I cancer, we compared the 37 HI with the 36 stage I NSCLC patients (out of 56 in total). EMAD combined with RF revealed a remarkably high level of performance, achieving ACC 0.99, AUC 1.00, and Se 1.00, at an imposed Sp of 0.95, with 4 and 3 bp end motifs (Fig. 5A). These results were supported by PCA 2D-projection, which showed the formation of distinct clusters for HI and stage I NSCLC (Fig. 5B), and for chosen end motifs (Fig. 5C) using 3 bp motifs. By contrast, the use of EMAD with RF to discriminate between different NSCLC stages was unsuccessful, with nearly 50% of the samples being predicted at the incorrect stage (Fig. 5D). Se, consequently, was close to zero. This result is in stark contrast to the near-perfect distinction made between HI *and* pooled stage I, II, and III NSCLC (Fig. 3A).

**Figure 5.**
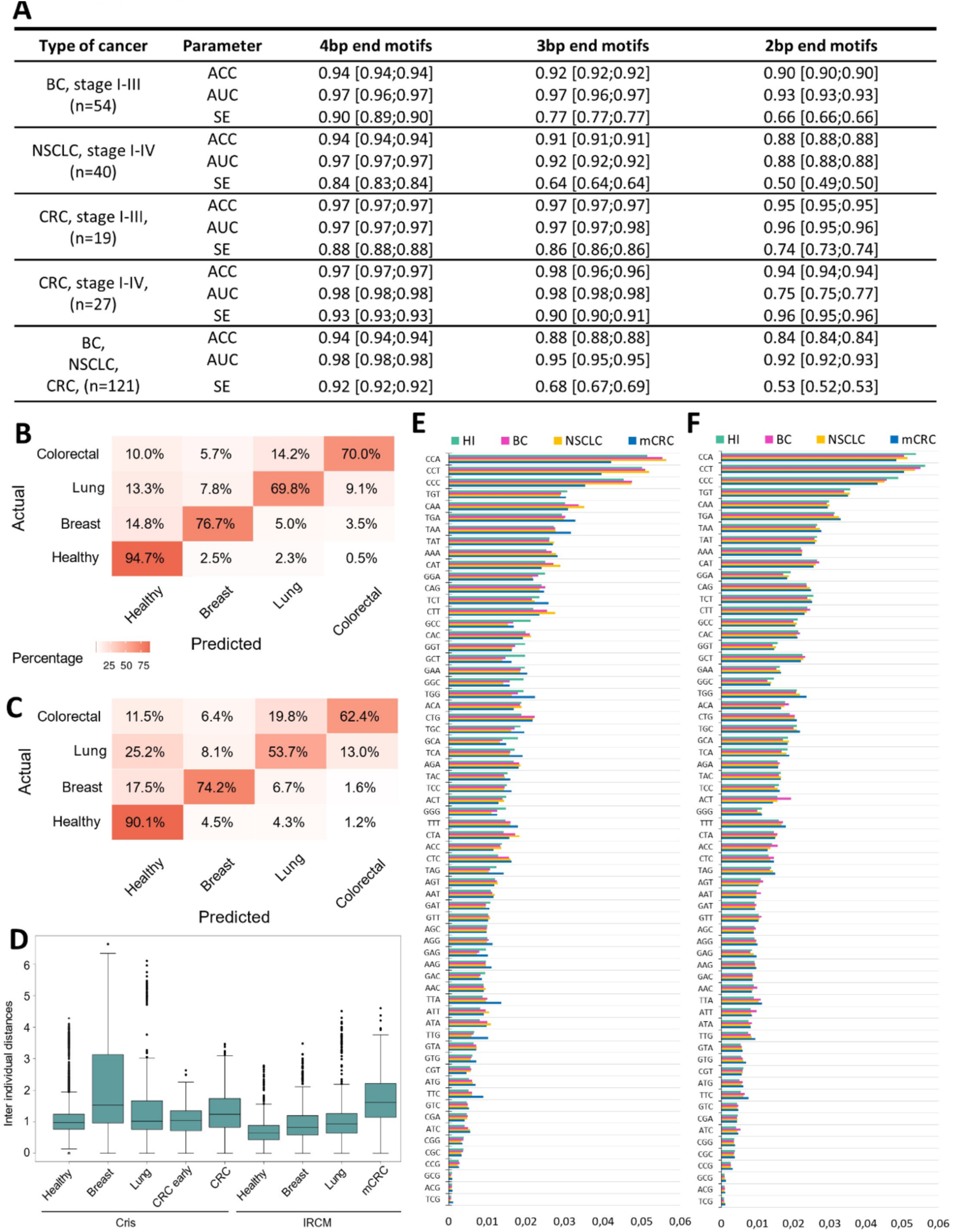
EMAD validation using external data. **(A)** EMAD with Random Forest (RF) model performance for detection of breast cancer (BC), non-small lung cancer (NSCLC), colorectal cancer (CRC), and a pooled group of all three cancers, versus healthy individuals (HI). Models were evaluated at 0.95 Sp. **(B, C)** Confusion matrices showing the classification performance of the RF model in distinguishing breast, lung, colon cancer, and healthy samples based on 4 bp **(B)** and 3 bp **(C)** end motif relative frequencies. The featured values represent the average of 100 times repeated 5-fold cross-validation confusion matrices on the test data. **(D)** Distribution of inter-individual distances for healthy individuals, breast, lung and colorectal cancer (CRC, mCRC for metastatic CRC) for our (IRCM) and Cristiano cohorts using 3 bp end motif relative frequencies. **(E,F)** Relative frequencies of 3 bp end motifs for HI, BC, NSCLC and mCRC in our dataset **(E)** and the external dataset of Cristiano et al. **(F)**.

### Confirmation with publicly available cohorts

To confirm EMAD’s potential, we exploited independent cirDNA data from the Cristiano *et al.* study ^10^, which we retrieved from FinaleDB ^31^. This dataset included 245 HI, 54 BC, 40 NSCLC, and 27 CRC samples. Among the CRC samples, 19 were stage I-III and 8 stage IV (mCRC). CirDNA fragments were sequenced using a DSP library preparation protocol similar to ours (NEB Next kit followed by Illumina paired-end sequencing). We applied EMAD and RF as above, with the performance being assessed by 5-fold CV. The results remained close to those we obtained from our data using 4 bp end motifs (Fig. 6A). However, the performance here decreased when 3 and 2 bp end motifs were used. Pooling all three cancers, as we did above, decreased performance further with 3 bp, and even more so with 2 bp, while it remained close to individual cohorts using 4 bp motifs. In the task of distinguishing between cancer type, we achieved an accuracy of classification > 0.70 with 4 bp (Fig. 6B) and > 0.55 with 3 bp (Fig. 6C).

**Figure 6.**
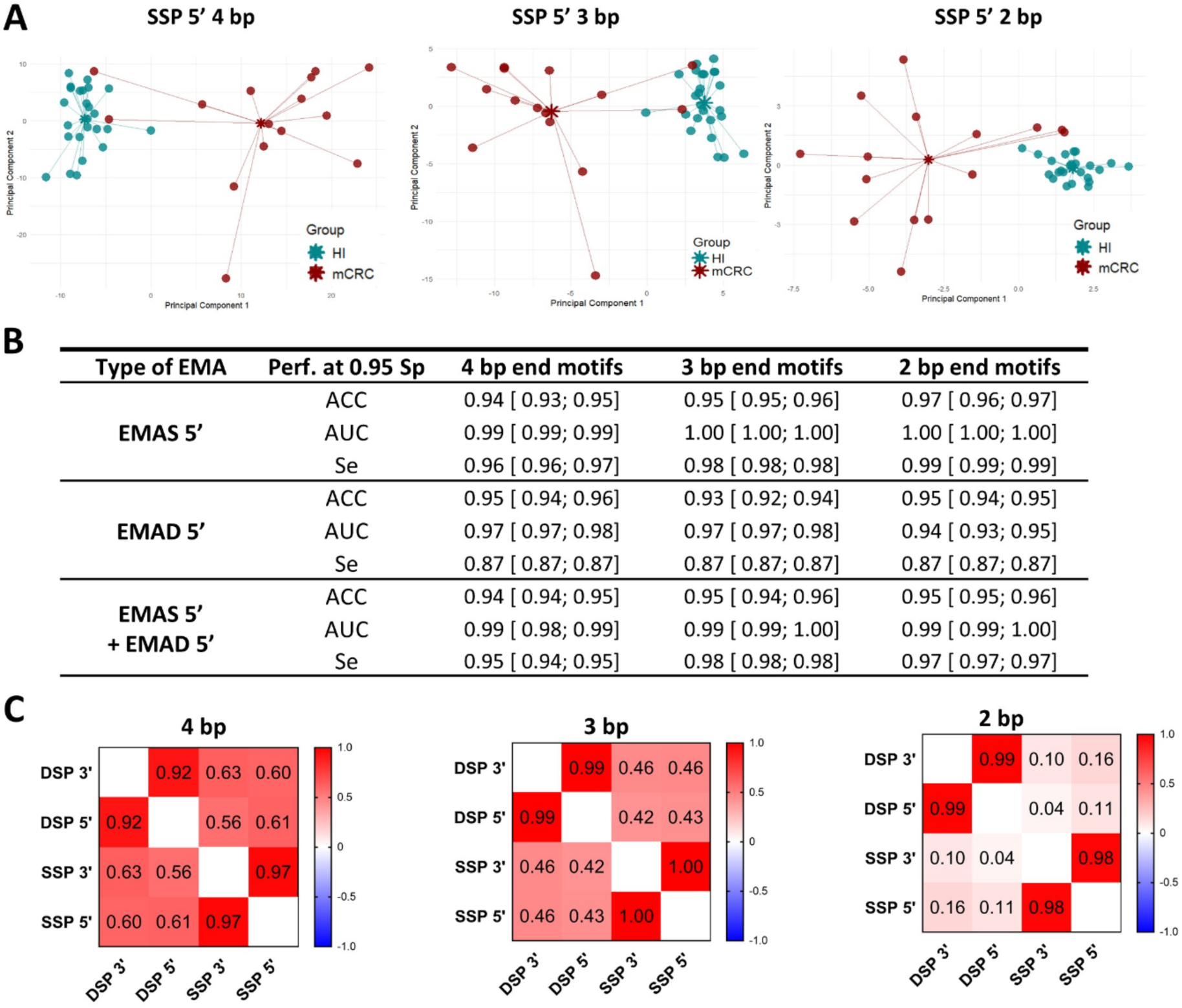
EMAD and EMAS comparison on mCRC. **(A)** PCA 2D-projections of 4, 3, and 2 bp end motif relative frequencies in SSP-sWGS and DSP-sWGS data (25 healthy individuals (HI) and 15 patients with metastatic colorectal cancer (mCRC)). **(B)** Performance of EMAS and EMAD with Random Forest (RF) for the detection of mCRC versus HI. Performance statistics were obtained imposing 0.95 sensitivity. Note that the HI involved in this analysis only included those that were sequenced both in SSP-sWGS and DSP-wWGS (25 out of the 37 HI used in Fig. 2), hence the slight differences in performance statistics. **(C)** Correlation matrices for mCRC displaying the relationships between relative frequency of cirDNA fragment end motifs across different library preparation methods.

While this use of EMAD with Cristiano *et al.* data for the task of cancer detection produced only a slightly reduced level of performance, the loss of discriminatory power was greater in the more difficult task of distinguishing cancer types. This suggests that the Cristiano *et al.* ^10^ data were more difficult to model with an EMA approach. Indeed, comparing all the distances between end motif relative frequencies of distinct individuals within a given cohort, we found larger values within all Cristiano *et al.* cohorts except for mCRC (Fig. 6D). As expected, HI distances were smaller than within cancer cohorts. Heterogeneity of end motif frequencies within a cohort was not directly related to detection performance. For instance, the Cristiano *et al.* BC cohort was more heterogeneous than their NSCLC, but the performance levels for both cohorts was the same.

We compared the relative frequencies of the 3 bp end motifs in our and the Cristiano data, in the HI (Fig. 6E), BC, NSCLC, and CRC cohorts (Fig. 6F). Although globally similar, more attenuated differences were observed within the Cristiano samples, which was in line with the higher heterogeneity of those samples (Fig. 6D). We also observed variations in the 5’ and 3’ DSP end motif frequencies, which might indicate the presence of adaptor sequences which were not completely cleaned (compare with Fig. 6F). Furthermore, for a given end motif, frequency variations *per* sample type were slightly different between the two data sources, which could be indicative of a logical platform bias (Fig. 6).

### Comparing EMAD and EMAS in mCRC plasma samples

To compare data obtained from SSP-sWGS and DSP-sWGS, we sequenced 25 HI and 15 mCRC patients twice, employing both DNA library preparation protocols (Fig. 1). EMA applied to SSP-sWGS was named EMAS to distinguish it from EMAD. As illustrated schematically in Fig. 1, the SSP and DSP protocols are expected to generate different end motif sequences, which in turn should impact their frequency distributions. This expectation was confirmed by our analysis: the same end motifs exhibited markedly different frequencies depending on the protocol used, both in HI (Fig. S3) and in mCRC (Fig. S4). Interestingly, 5’ and 3’ SSP end motif frequencies were as close as those in DSP, indicating an absence of orientation preference (Fig. S3). Accordingly, we decided to consider only 5’ end motifs in both DSP and SSP.

PCA 2D-projection of SSP end motif relative frequencies showed that HI tended to cluster together while mCRC patients were separated but more diffuse for 4, 3, and 2 bp motifs (Fig. 7A). Encouraged by this result, we again built RF models that we assessed with 5-fold CV considering EMAD and EMAS, and also the combinations of DSP and SSP data (Fig. 7B). The main observation which emerged from this was that EMAS yielded slightly better performance than EMAD. While 4 bp end motifs showed a slightly higher level of performance as compared to 3 or 2 bp when EMAD was used, EMAS exhibited higher performance level when examining 2 bp end motifs (0.95, 0.97, 0.87 and 0.97, 1.00, 0.99; ACC, AUC, Se, respectively). EMAS performance is greater than EMAD particularly due to a rather low sensitivity using EMAD (0.87 when specificity is 0.95). As anticipated, a strong Spearman correlation was found when comparing 5’ and 3’ end motifs relative frequencies at any length for either given protocol. By contrast, the discrepancy between EMAS and EMAD data was highlighted in the lack of association produced when 2 bp end motifs were used, in contrast to 4 bp end motifs (Fig. 7C). Note, when analyzing 3 bp end motifs the EMAS + EMAD data combination did not improve performance over EMAS alone. When analyzing mCRC samples, a large variation in the frequency of each end motif was observed between EMAD and EMAS (Fig. 4C and D; Fig. S3 and 4). For EMAS, the end motifs that showed the highest relative frequency were AAA, AGA, GGA, GAA, whereas for EMAD they were CCA, CCT, CCC and TGT (Fig. 4C and D). While the end motifs with the lowest relative frequency were identical for both protocols (TCG, ACG, CCG, GCG, CGG, CGA, CGC and CGT), they appeared to group altogether following EMAS (Fig. 4D). Regardless of whether EMAD or EMAS were used, the end motif frequency which showed the highest increase in mCRC samples were similar (TCG, TTC, TTG, and TTA). For EMAS and EMAD separately, the end motifs which showed the lowest relative frequency change were CCC, CCT, ACC, GCC, GCT and CGT, and CCC, CCT, CCA, GCC, GCT, and CGT, respectively (Fig. 4C and D)The top10 more elevated SHAP values using EMAS for mCRC GGT, GGC, GGA, GCC, GCA,, GGG, CAT, CAA,CTT and TCG, while AAA, AGA, AGG, GAA,GGA, TGT,TGA,TGG showed the highest level of relative frequency (Fig. S5). Note, GGG appears as the only one common end motif with high SHAP value for BC, NSCLC and mCRC as well as for SSP. The comparison of the top SHAP values vs the top relative frequencies illustrated that there is unexpectedly a poor direct relationship between sequencing and Random Forest data when comparing EMAS to EMAD, or EMAD in the three examined cancer types. Consequently, the method provides inventive information (in addition to indicate the end motifs that provide optimal distinction between cancer and healthy individuals), to indicate some end motifs showing specificity to the use of SSP vs DSP sWGS.

**Figure 7.**
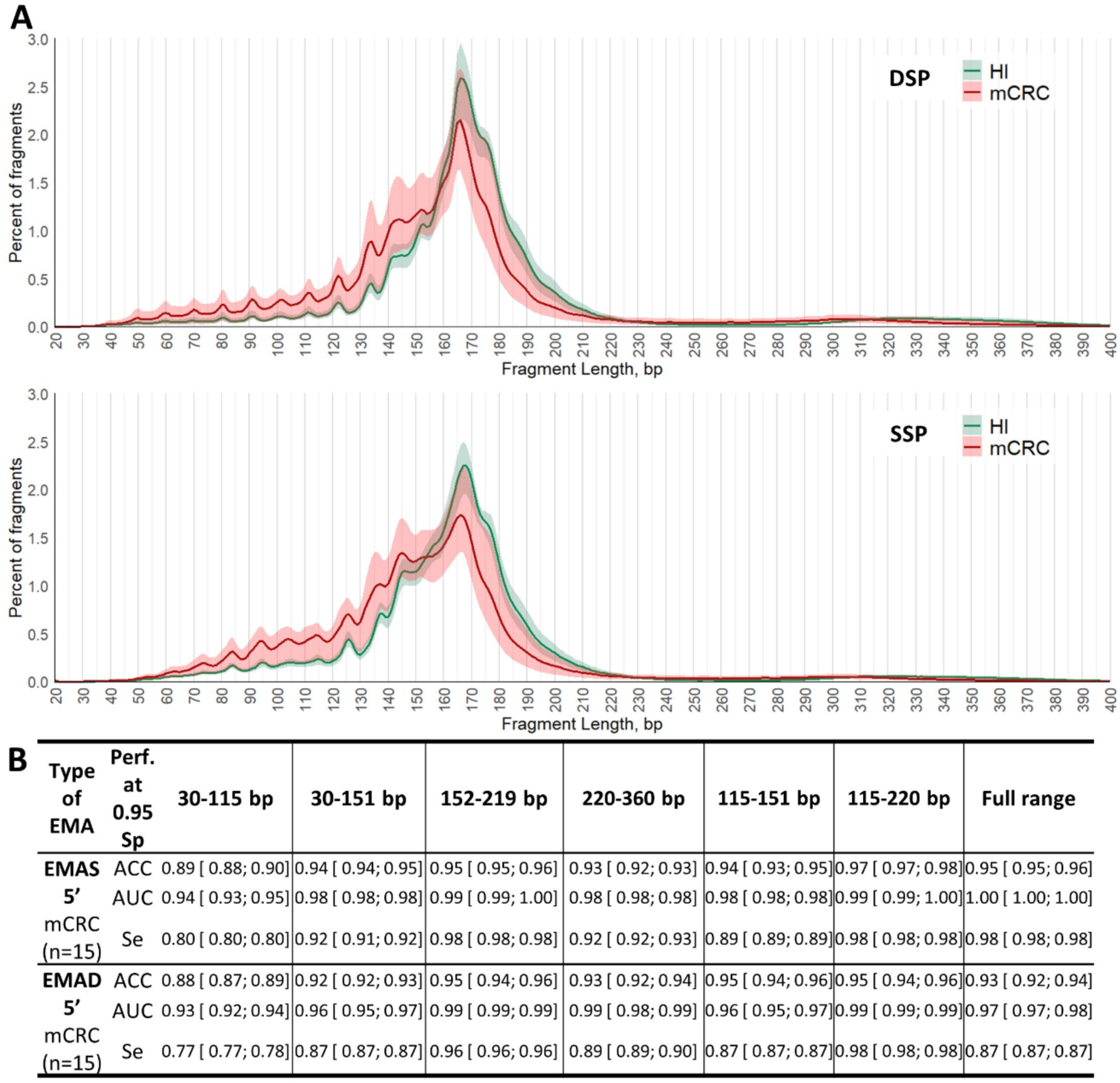
Analysis of cirDNA end motifs in relation with fragments sizes in DSP and SSP libraries. **(A)** Size profile of cirDNA fragments obtained by SSP or DSP library preparation methods from 25 HI versus 15 mCRC patients (mean with 95%CI). **(B)** Performance of EMAS and EMAD with Random Forest models for the detection of mCRC. A specificity of 0.95 was imposed, 5-fold cross-validation repeated 100 times.

### EMAD and EMAS efficiency with respect to cirDNA fragment size

The size profile of cirDNA fragments is known to differ between HI and cancer patients, as illustrated by Fig. 8A, which features DSP-sWGS and SSP-sWGS data for both HI and mCRC patients. As compared to HI, the DSP-sWGS and SSP-sWGS cirDNA fragment size profile of mCRC shows a shift towards shorter values, while also revealing a chromatosome-associated major peak at 167 bp, as previously described ^17^. In addition, the SSP-sWGS size profile exhibited a higher proportion of fragments below ∼152 bp, as compared to the DSP-sWGS size profile. It should be noted that HI and mCRC size profile curves crossed each other at 152 bp and 220 bp, revealing a higher proportion of fragments in the 30 - 153 bp and 220 - 360 bp size ranges in mCRC patients, irrespective of the use of SSP-or DSP-sWGS.

To assess the potential existence of cirDNA fragment size ranges more favorable to EMAD or EMAS, we considered specific size ranges, *i.e.*, 30-115 bp, 30 - 151 bp, 115-151 bp, 115-220 bp, 152 - 219 bp, and 220 - 360 bp (Fig. 8B). The analysis was limited to 3 bp motifs for simplicity. We found that, restricting analysis to 115-220 bp and 152 - 219 bp size range produced the highest level of performance for EMAD. EMAS performance was optimal when analyzing 115-220 bp and 152 - 219 bp size range but also the size full range and the 220-360 bp size range. Clearly data showed that EMAS outperformed EMAD for 30-150 bp, 115-151 bp and 220-360 bp as well as for the full range. In agreement with the results in Fig. 7B, EMAS globally delivered better performance levels than EMAD.

## DISCUSSION

We introduced a generic approach to cancer detection and classification based on the statistical modeling of individual sample cirDNA fragment end motif relative frequencies obtained after sWGS (EMA). We considered both DSP and SSP DNA library preparation strategies, and we named the corresponding end motif analysis methods EMAD and EMAS, respectively. The choice of the specific statistical model is free in this context. RF models were used to evaluate EMA for cancer detection. Starting with the most common sample preparation strategy, *i.e.*, DSP, we found EMAD to be extremely good at discriminating between healthy individuals and individuals affected by cancer. For instance, in the case of plasma from BC patients (stage II-III), EMAD Se and Sp were 1.00. EMAD achieved similar performance levels with stage I NSCLC patients. EMAD performance with mCRC patients remained very high, albeit slightly reduced, with a Se of 0.87. Early detection of noninvasive lung cancer was investigated by Chabon *et al*., whose method exploited cirDNA mutations and fragment sizes ^3^ achieving Se 0.41 at Sp 0.98, or Se 0.63 at Sp 0.80 for stage I NSCLC, respectively. The same work reported Se 0.69 and 0.75 for stage II and III NSCLC, respectively, at Sp 0.80. Wong *et al.*, who studied the early detection of Li-Fraumeni syndrome by means of cirDNA ^32^ combined copy number variation, mutation, and fragment size information to obtain positive and negative predictive values of 0.54 and 0.95, respectively. Bao *et al*., used 6 bp end motifs along with other cirDNA fragments features (fragment sized, breakpoint motifs, copy number variation), they achieved Se 0.96 at Sp 0.95 when comparing HI with a pool of primary liver cancer (PLC, stage I-III), CRC (stages 0 and I), and lung adenocarcinoma (LUAD, all stage I) patients ^33^. Even more specifically, our results should be compared with approaches which, like ours, solely involved end motifs. For instance, Jiang *et al.* exploited 4 bp 5’ end motifs to detect hepatocellular carcinoma, achieving AUC 0.89 ^23^. EMIT, relying on large language models (transformers) applied to 4 bp end motifs, obtained performance levels similar to ours ^34^. More precisely, EMIT achieved slightly lower AUC with 2 megabytes of model variables, and achieved AUC close to 1 with 32 megabytes. In other, more recent studies also worthy of comparison, Shen *et al*. ^34^ and Zhou *et al*. ^24^ described end motif analysis based on deep learning models, producing similar performance levels to ours. Elsewhere, in combining fragment sizes, genomic coverage, and end motifs towards detection of lung cancer, Lee *et al.* achieved AUC 0.94 ^35^. Comparisons of the above results with our own suggest that the intrinsic information present in end motifs is readily accessible to off-the-shelf ML. Given that we used Cristiano *et al.* data as an external dataset, it was interesting to compare our results with their seminal work ^10^. Their method, named DELFI, relied on cirDNA fragment size distributions in genomic windows. DELFI achieved Se 0.70 and 0.81 in BC and CRC *versus* HI, respectively, at 0.95 specificity. By comparison, the application of EMAD to the same data obtained Se 0.86 in both cases. The NSCLC cohort in Cristiano *et al*’s original publication was smaller than what was publicly released and used as external data by us (12 out of 40 patients). Consequently, we did not compare NSCLC performance.

We showed that EMAD has the potential to retain cancer type-specific signatures. In particular, we exploited this feature to build automatic classifiers capable of classifying samples according to whether they are HI, BC, mCRC, or NSCLC, and to do so with a level of accuracy superior to 93% with 4 bp end motifs. By comparison, Bao *et al*, exploiting 6 bp end motifs to consider lung, CRC, and liver cancers, were able to predict cancer origin with an accuracy of 91% ^33^, which is comparable to our performance. Similarly, DELFI managed to identify tumor types among BC, NSCLC, CRC, ovary, pancreatic, bile duct, and gastric cancer patients with an overall accuracy of 61% ^10^. Also, another method named CRAG, which exploits the concept of fragmentation genomic hotspots in order to detect cancer, was applied to Cristiano *et al.* data. This predicted cancer tissue of origin with an accuracy of 0.67 for pancreatic cancer, and 0.97 for BC ^36^.

Remarkably, we discovered that very short 2 bp end motifs could be used to distinguish HI from cancer patients. In the more challenging task of distinguishing between different cancer types, 2 bp end motifs performed more modestly, although the accuracy of classification remained nonetheless superior to 84%. Switching to 3 bp raised this accuracy to 91%, while the use of 4 bp motifs achieved an accuracy in cancer type classification > 93%. These results indicate that the first two positions of the fragment sequence have the highest discriminatory power. This could be due to nuclease preferences or variations in cirDNA fragment stability. Because we observed cancer-specific changes in end motif relative frequencies, we are inclined to believe that these observations in fact reflect a combination of nuclease preferences with variations in particular nuclease activities, depending on the disease ^23^. To this particular point, it should be noted that Lo’s group revealed that distinct nucleases are associated with different last nucleotide biases, namely: (i), deoxyribonuclease 1L3 (DNASE1L3) nicks DNA in a manner that leads to a preponderance of C ends; (ii), deoxyribonuclease 1 (DNASE1) preferentially cuts DNA into T end (32); and (iii), DNA fragmentation factor subunit beta (DFFB) plays an important role in the fragmentation of DNA *de novo* released from dying cells ^14^.

Regarding end motif frequency, it should first be noted that the CCA, CCT, and CCC motifs are by far the most frequent in both healthy individuals and cancer patients, as previously observed ^23,37–39^. The TCG, ACG, and GCG motifs are the least frequent in both healthy individuals and cancer patients, as previously observed by Moldovan *et al*. ^39^. Previous reports exactly ^39^ or partially ^34,38,40^ described those same motifs that show the highest relative frequency difference, as well as those that show a reduced relative frequency difference in cancer patients.

The following data from DSP sequencing indicates that: (i), the CCA motif is consistently the most frequent motif in the three cancer types studied, as previously described for instance by Jiang *et al.* ^23,41^, Jin *et al.* ^38^, and Moldovan *et al.* ^39^; (ii), the motifs with the highest frequency and the greatest difference in relative frequency compared to healthy individuals are identical, regardless of the cancer type studied; (iii), the motifs with the lowest relative frequency are the same for all three cancer types; and (iv), both the most and the least frequent motifs we found when examining our cohorts of patients with BC, NSCLC, and mCRC are exactly the same as those found from the external dataset of Cristiano *et al*. ^10^. Altogether these four points suggest at the most a moderate influence of a possible bias due to the different pre-analytical conditions or DNA library preparation kits used in our and above indicated cohorts. Note, while we used the same sequencing DNA library kit in our study, the pre-analytical conditions largely differed among the BC, NSCLC and mCRC cohort samples. For instance, for blood collection EDTA tubes were used for HI, citrate tubes for BC, and Streck cell preservation tubes for the mCRC and NSCLC samples. An EMA prospective analysis of samples from an *ad hoc* cohort of lung and colon cancers, subjected to exactly the same preanalytical conditions, is currently being conducted, to estimate the importance of sampling bias due to pre-analytical conditions. This topic has never before been accurately explored in the literature. While the relative frequency pattern appears similar for stage II-III BC and stage I-III NSCLC, this is not entirely true for mCRC. It’s worth noting that, despite this, samples of the BC and NSCLC types could nonetheless be distinguished by EMAD.

It should be noted that Moldovan *et al.* ^39^ found that, in CRC, the end motifs which are most depleted correspond to those also detected in our study. In addition, some of the motifs this study identified as having the lowest relative frequency are identical to those we identified (GCG, CCG, CGA). Furthermore, our results show that the three motifs with the lowest relative frequency show a positive difference in mCRC, while the difference is negative in BC and NSCLC samples. Another notable point of contrast concerns the three most frequent end motifs which show a negative relative frequency change in CRC patients and a positive relative frequency change in BC and NSCLC patients. This would appear to suggest that there is a variation in the pattern of distribution of motifs between mCRC on the one hand, and BC and NSCLC on the other hand. Although investigating the relative frequency change of each individual end motif is informative as to mechanisms involved in fragmentation, the individual contribution of any particular end motif to the discrimination between the cirDNA of cancer patients and of HI, as performed by Random Forest classification, derives from both the value of the relative frequency change and the level of variation of that end motif’s relative frequency values.

The distribution of relative frequencies of end motifs that we obtained from DSP-sWGS in samples from HI is consistent with the general observation described by Noé *et al.* ^22^. That observation indicates that the relative frequency is enriched by specific motifs, in particular those with a thymine or an adenine before the beginning of the fragment and followed by two cytosines, and those with a cytosine followed by a guanine. Our results correspond overwhelmingly to the former group and rarely to the latter. Zhou *et al*. ^24^ recently defined different “founder” profiles related to a group of specific motifs linked to a type of fragmentation. One of those founder profiles concerns the activity of DNase1L3, which generally leads to the production of motifs starting with CC. As indicated by Zhou *et al*. ^24^, such end motif profiles would be useful to inform physiological conditions such as autoimmune diseases (Systemic Lupus Erythematosus, SLE) and cancer. SLE is strongly associated with a high concentration of cirDNA and the production of neutrophil extracellular traps (NETs). We have previously shown the association of NETs with cirDNA production in mCRC ^42^ and stage III colon cancer 43, particularly post-surgery. More specifically, we have demonstrated that NETs are a significant cause of the cirDNA release, and that the size distribution profile of cirDNA fragments is specific to NETs degradation, not only in CRC, but also in patients with SLE ^44^. These findings suggest that the mechanisms that generate NETs production may be linked to a specific pattern profile in the case of SLE or cancer. Furthermore, we demonstrated that enzymes physically associated with NETs, such as myeloperoxidase and neutrophil elastase, contribute to the degradation of NETs, and could be at the origin of the founding profile IV defined by Zhou *et al*. ^24^. This founder profile represents a nonspecific cleavage pattern, and its examination also appeared very effective to differentiate control individuals and patients with cancer. The impact of NETs on end motif frequency is under active investigation by our team.

This study is the very first to compare the use of commonly applied DSP protocol with the rarely applied SSP protocol, with respect to cirDNA end motif analysis. DSP is a commonly used straightforward protocol which can be completed in just a few hours, while SSP typically takes much longer ^29^. Despite that drawback, SSP has a number of specific advantages over DSP. Indeed, it was specifically developed to overcome the limitations of conventional DNA library methods, namely to recover damaged and short double-stranded DNA fragments, to facilitate forensic and ancient DNA analysis ^45^. The analytical mechanisms which differentiate SSP from DSP, and the phenomena they exploit, are especially relevant for end motif analysis. Native cirDNA fragments have two DNA strands, which vary in sequence and coordinates due to the presence of 5’ or 3’ overhangs. In the DSP protocol, an end repair step completes the 5’ overhangs and polishes the 3’ overhangs, making them the same length (blunt ends). By contrast, SSP does not include any end repair step, and therefore should reveal end motifs that constitute more accurate reporters of actual nuclease activity. Our innovative EMA method not only takes these differences into account, but indeed exploits them. To do so, first we demonstrated that EMAS achieved high performance levels comparable to EMAD in cancer detection. Second, we explored the differences in 5’ and 3’ end motifs. As expected, DSP end motifs at the 5’ and 3’ fragment extremities harbored almost identical relative frequencies. This was also the case for SSP end motifs, which we interpret as the absence of the orientational preference of the enzymes involved. Except for this particular aspect, DSP and SSP end motif frequencies were very different. To the best of our knowledge, these differences have never been described before in the literature. Overall, when comparing SSP-and DSP-sWGS data, the frequency ranking differed significantly between the two, in contrast to the relative frequency change ranking, which did not. Nevertheless, CCA, CCT, and CCC frequencies were considerably reduced when SSP was used. Specifically, the highly studied CCA end motif showed high variation in both frequency and relative frequency change, being more or less ubiquitous among all end motifs following SSP and DSP. However, our EMAS data does not strictly conform to the expected patterns of individual end motif frequency as defined either by Noé *et al*. ^22^ or Zhou *et al*. ^24^. Consequently, the use of SSP provides different data potentially enlarging the efficiency of cirDNA fragment end motif analysis.

Lastly, we explored the relation between end motif relative frequencies and cirDNA fragment sizes. Considering the commonly observed differences between the cirDNA fragment size ranges of healthy individuals and individuals with cancer, we defined three fragment size ranges: 30 - 151 bp, 115-220 bp, 152 - 219 bp, and 220 - 360 bp. Data analysis revealed a potential advantage for EMAD in working with 115 - 219 bp fragments, as opposed to exploiting the complete size range. In the case of EMAS, the same 115 - 219 bp size range delivered performance levels identical to the complete size range, and in addition, EMAS showed higher performance as compared to EMAD in each of the shorter or longer size ranges. To assess the potential existence of cirDNA fragment size ranges more favorable to EMAD or EMAS, we considered specific size ranges, *i.e.*, 30-115 bp, 30 - 151 bp, 115-151 bp, 115-219 bp, 152 - 219 bp, and 220 - 360 bp. The analysis was limited to 3 bp motifs for simplicity. We found that, restricting analysis to 115-220 bp and 152 - 219 bp size range produced the highest level of performance for EMAD. EMAS performance was optimal when analyzing 115-220 bp and 152 - 219 bp size range but also the size full range and the 220-360 bp size range. Clearly data showed that EMAS outperformed EMAD for 30-150 bp, 115-151 bp and 220-360 bp as well as for the full range. In agreement with the results in Fig. 8B, EMAS globally delivered better performance levels than EMAD.

This study has several limitations. Further validation with a larger number of individual samples is required. In addition, the study was conducted retrospectively and consequently; that is to say, first, that it does not fully reflect the real-world context of cancer screening; and, second, it does not provide rigorous standardization, especially with regard to the evaluation of possible bias due to pre-analytical conditions. Since they were not so far studied and not reported in the literature, a well-designed prospective study is necessary. These limitations are at this time undertaken in ongoing clinical studies using EMAS in different cancers.

Our study highlights the possibilities of refining cirDNA end motif analysis, as well as extending its versatility, by considering a number of variables, namely: motif size, fragment size range, the use of double-or single-stranded DNA library preparations, and end motif frequency variation, in both cancer patients and healthy individuals. In this way, our work represents a significant advance in the optimization of end motif analysis of cirDNA fragments towards cancer detection. Indeed, to that end, our observations specifically lead us to recommend the use of sWGS, and reveal the benefits in using SSP-over DSP-sWGS. Overall, there are clear discrepancies between data obtained from EMAD and from EMAS: (i), EMAS’s performance levels were higher than EMAD’s; (ii), EMAD produced higher performance levels when 4 bp end motif size were examined, while EMAS performance levels appeared to be less dependent on end motif size; (iii), the correlation between EMAD and EMAS data appeared to moderate at best; (iv), their association decreased as end motif size decreased, to being null when 2 bp end motifs were examined; (v), the relative frequency from EMAS of each end motif is different of that obtained from EMAD; and (vi), the performance of EMAS was optimal when the size full range was examined, while EMAD’s performance was optimal for the 115 - 220 bp range. This enabled to optimize cirDNA end motif analysis by combining EMAS on the full range of fragment sizes, and analysis of three base pairs end motifs.

Overall, this work enabled us to considerably deepen our understanding of cirDNA fragmentomics, and to optimize this approach towards cancer detection from revealing significant features of end motifs which were previously unobserved. EMAS performance of cirDNA end motif analysis in detecting cancer and distinguishing cancer types that combines cost-effective sWGS, SSP, and ML assistance would warrant future large-scale deployment of this approach in diverse patient populations.

## MATERIALS AND METHODS

### Healthy individuals and patients

We analyzed cirDNA from the plasma of 37 HI (Supplementary Table S1) obtained from the Etablissement Français du Sang (EFS), which is Montpellier’s blood transfusion center, under the EFS-PM N° PLER2024-003-R agreement number. These samples were initially screened (virology, serology, immunology, blood count), and those showing any abnormality were ruled out.

We analyzed cirDNA from the plasma of 50 patients with stage II-III breast cancer (Supplementary Table S1) from the Centre Léon Bérard, which is one of the UNICANCER GastroIntestinal Group (UCGI), Paris, France. Samples were managed by the Biological Resource Center (BRC) of the Centre Léon Bérard (no. CRB-CLB BB-0033-00050). This was done within the framework of the MyProbe project (an RHU program of the French National Research Agency grant ANR-17-RHUS-008). The patients included had HR+/HER-tumors, and were treated by tumor surgical resection. Blood was collected on the day of surgery before the administration of anesthesia. No patients had received neoadjuvant treatment before blood collection.

We also analyzed cirDNA from 56 patients with stage I-III NSCLC, recruited within the prospective observational clinical trial LUNGDOC (NCT05165160) at Montpellier University Hospital (Supplementary Table S1). The patients included underwent surgical tumor resection, and blood was collected preoperatively.

Lastly, we analyzed cirDNA from 15 patients with stage IV mCRC from the screening procedure of the UCGI 28 PANIRINOX study (NCT02980510/ EUCT 2024-510645-34-00). PANIRINOX sponsored by UNICANCER is an interventional study using cirDNA as a companion test for the selection of mCRC patients towards anti-EGFR targeted therapy, using the IntPlex® method ^15,46^. Eligible patients were recruited in the PANIRINOX screening procedures at diagnosis. Included patients had a histologically confirmed colorectal adenocarcinoma, an untreated synchronous, or metachronous metastatic disease deemed unresectable with curative intent.

Written informed consent was obtained from all participants in all cohorts.

### Blood collection and plasma isolation

Samples were handled according to pre-analytical guidelines previously established by our group ^47^. HI blood were collected in 6 mL EDTA tubes (K2E, REF 367525, BD Vacutainer) or 10 mL STRECK tubes (Cell-Free DNA BCT CE, n4015, STRECK), in 2.7 mL citrate tubes (9NC 0.109M, REF 363048, BD Vacutainer) for BC patients, and in 10 mL STRECK tubes for NSCLC and mCRC patients. Using our preanalytical procedure, we found that tube type had no impact on cirDNA fragmentation (data not shown). Blood was then centrifuged at 1,200 g at 4 °C for 10 min. Supernatants were isolated in sterile 1.5 mL Eppendorf tubes and stored at – 20 °C (freezing was done for HI, BC and mCRC, whereas NSCLC samples were processed without plasma freezing after 1^st^ centrifugation). Plasma samples were then centrifuged at 16,000 g at 4 °C for 10 min. Plasma was distributed by 1.1 mL to the 1.5 mL Eppendorf tubes. Afterward, plasma was stored at –20 °C until cirDNA extraction. Hemolytic, icteric or turbid (triglycerides) plasma samples were discarded.

### External dataset

We downloaded a public data of HI, NSCLC, BC, and CRC individuals ^10^ from FinaleDB ^31^. There were 245 HI samples (ignoring technical replicates available for 16 of these HI), 79 NSCLC samples out of which we only kept the 40 samples corresponding to pre-treatment and preoperative treatment naïve conditions, 27 CRC samples, and 54 BC samples.

### DNA extraction from plasma

CirDNA was extracted from 1 mL of plasma using the QIAamp DNA Mini Blood kit (Qiagen, Cat. No.51104) following the ‘‘Blood and body fluid protocol’’, using 80 μL of elution volume ^15,47^. DNA extracts were kept at – 20 °C until used.

### DNA library preparation for sWGS

DSP (double-strand protocol) and SSP (single-strand protocol) libraries were prepared. SSP allows the integration of single-and double-stranded DNA in the library. DSP libraries were prepared with the NEB Next® Ultra™ II kit. SSP libraries were prepared with the Swift ACCEL-NGS® 1S PLUS kit. For both preparations, a minimum of 2 ng of cirDNA was engaged without fragmentation, and each kit provider’s recommendations were followed. These recommendations can be summarized as follows. For DSP (with the NEB kit), Illumina paired-end adaptor oligonucleotides are ligated on repaired A-tailed fragments, then purified by solid-phase reversible immobilization (SPRI) and enriched by 11 PCR cycles with UDI primers indexing, then SPRI purified again. For SSP (with the Swift kit), heat denaturation is performed first, to allow the conversion of all DNA into single strands; this protocol allows, simultaneously, the extension of single-stranded fragments and the attachment of adapters to the end of these fragments. Adaptase simultaneously links an adapter to and extends their 3’ end. The complementary strand is then synthesized by a primer extension. After SPRI purification, a second adapter is ligated at the other end, purified by SPRI, and the product enriched by 11 cycles of PCR, then purified by SPRI again. For both types of preparation, SPRI purification was adjusted to keep the small fragments around 70 bp of insert. Finally, the libraries to be sequenced were precisely quantified by Q-PCR, in order to load the appropriate DNA quantity to the Illumina sequencer, and to obtain a minimum of 1.5 million clusters.

### Fragment size and end motifs extraction from sWGS data

All libraries were sequenced on MiSeq 500 or NovaSeq (Illumina) as Paired-end 100 bp reads. Image analysis and base calling were performed using Illumina Real Time Analysis with default parameters. The individual barcoded paired-end reads were trimmed with Cutadapt v1.10, to remove the adapters and discard trimmed reads shorter than 20 pb. Trimmed FASTQ files were aligned to the human reference genome (GRCH38), using the MEM algorithm in Burrows-Wheeler Aligner v0.7.15. Supplementary alignments, PCR duplicates, and reads with MapQ < 30 were filtered out from the analysis. Coverage values (mean±sd) were assessed for both the DSP and SSP for all sample groups. For the DSP, the mean coverage was 0.52 ± 0.24 in healthy individuals, 0.55 ± 0.13 in breast cancer (BC), 0.52 ± 0.08 in non-small cell lung cancer (NSCLC), and 0.67 ± 0.12 in metastatic colorectal cancer (mCRC). For the SSP, the mean coverage was 0.44 ± 0.07 in healthy individuals and 0.50 ± 0.09 in mCRC. The insert sizes (=fragment sizes) were then extracted from the aligned bam files with the TLEN column for all pairs of reads having an insert size between 0 and 1,000 bp. The frequency of each fragment size was then normalized to the total fragment count to generate relative size distributions.

We determined the end motif sequence of the cirDNA fragments based on the filtered BAM files aligned against the reference human genome GRCh38. End motifs were determined for 5’ and 3’ DNA fragment ends. The sequences of end motifs were extracted and compiled using the pysam and pandas Python packages. For fragments mapped on the positive strand, 5’ end motifs were extracted as the n first nucleotides, and 3’ end motifs as the n last nucleotides (n being the length of the end motifs analyzed, i.e., 3, 2 or 1 bp). For fragments mapped on the negative strand, 5’ end motifs were extracted as the reverse complement of the n first nucleotides, and 3’ end motifs as the reverse complement of n last nucleotides (n being the length of the end motifs analyzed, i.e., 3, 2 or 1 bp). The relative frequency of each motif within each individual sample (or the fragment size fraction of the sample) was computed. When comparing 5’ and 3’ end motifs frequencies, the reverse complement of 5’ end motifs were used.

### Fragment size and end motifs extraction from Cristiano data

Files downloaded from FinaleDB for the Cristiano data set contain the position of cirDNA fragments mapped on the human genome GRCh38 in a BED-like tab-separated value format. Fragments with a quality score < 30 were filtered out. BED-like files were converted into genuine BED format using the pybedtools package, and sequences for all fragments were retrieved from the reference genome. End motifs were determined for 5’ and 3’ DNA fragment ends using the same approach as described above for sWGS data from our cohorts.

### Machine learning

ML was performed using R (v 4.3.3) and the software packages: randomForest (v 4.7-1.2) for RF models, caret (v 7.0-1) for cross-validation and confusion matrix extraction, pROC (v 1.18.5) for AUC calculation, MASS (v 7.3-60.0.1) and plotly (v 4.10.4) for LDA, ggplot2 (3.5.1) for plotting, and built-in R function prcomp() for PCA. We performed 5-fold CV of the RF models (100 repetitions). Reported confusion matrices represent the average of 100 such matrices obtained on the test sets of 100 repeated 5-fold CV. For the Cristiano data set, the imbalance of the healthy and cancer classes distribution was corrected using SMOTE.

### SHAP

To interpret the contributions of individual features to the predictions of the Random Forest classifier, we employed SHAP (SHapley Additive exPlanations) values. SHAP is a game-theoretic approach that attributes a model’s output to its input features based on Shapley values, providing both local and global interpretability^48^. We implemented SHAP analysis using the iml R package (version 0.11.4), which supports model-agnostic explanations^49^.

The model was first trained using the Random Forest algorithm via the caret package without cross-validation and with 500 trees. The Predictor object from iml was used to wrap the fitted model, specifying both the feature matrix and class labels. SHAP values were then computed individually for each observation using the “Shapley$new()” function.

To summarize feature importance across all samples, we calculated the mean absolute SHAP value for each feature. This provided a global measure of how influential each variable was in the model’s decision-making process. Visualization of SHAP values, including feature importance plots, was performed using ggplot2.

## Statistical tests

For comparison of the end motif relative frequencies of HI and patients with cancer, we used the Wilcoxon rank sum test. When multiple hypotheses were tested, P-values were adjusted using the Benjamini-Hochberg (BH) procedure.

## Ethics statement

All participants provided written informed consent. The study was approved by local ethics committees and conducted in accordance with the Declaration of Helsinki. Healthy donor samples were obtained under EFS agreement PLER2024-003-R. Breast cancer samples were managed under the MyProbe RHU program (ANR-17-RHUS-008). NSCLC samples were from the LUNGDOC study (NCT05165160) and metastatic colorectal cancer samples from the PANIRINOX study (NCT02980510/EUCT 2024-510645-34-00).

## List of Supplementary Materials

Fig. S1 to S5

Table S1.

## Acknowledgements

This work was supported by the ANR RHU MYPROBE grant on “Molecular assays to predict outcome in early Breast cancer”, the UCGI 28 PANIRINOX trial entitled “a randomized phase II study assessing Panitumumab + FOLFIRINOX or mFOLFOX6 in RAS and BRAF wild type metastatic colorectal cancer patients (mCRC) selected from circulating DNA analysis”, the ANR RHU REVEAL “Reshape the evaluation efficiency and accuracy of non-small cell Lung cancer”. This work was partially supported by SIRIC Montpellier Cancer Grant INCa_Inserm_DGOS_12553, by MSD AVENIR [MSD-THRuST grant and by the Société Française des Acides Nucléiques Circulants (SFAC).

The authors thank UGBDI, Centre Leon Berard, Lyon, France – Dr Christine Lasset and Melinda Teyssier – for having collected clinical data of the Centre Leon Berard’s patients. The authors thank A. Kudriavtsev and M. Chekroun for their helpful discussions and C. Sanchez, F. Frayssinoux and B. Pastor for technical assistance. The authors thank Cormac Mc Carthy (Mc Carthy Consultant, Montpellier) for English editing (financial compensation). We thank the healthy individuals from the EFS who participated in this study. We wish to thank all the patients, family members and staff from all the units that participated in the study.

## Funding

This work was supported by the following programs: 1. The RHU REVEAL project, funded by the French National Research Agency (ANR) under the “Programme d’Investissements d’Avenir” (PIA) as part of the Hospital-University Research in Health (RHU) initiative. 2. The MYPROBE project (ANR-17-RHUS-0008), funded by the French National Research Agency (ANR). 3. The PANIRINOX clinical trial (UCGI-28, NCT02980510), a phase II study conducted in metastatic colorectal cancer, in which circulating cell-free DNA was used as a companion biomarker for therapeutic decision-making.

## Author contributions

Conceptualization: EP, MR, JC and ART. Data curation: EP, TM, BR, BB, FA, WJ,, STE, TB and MY. Formal Analysis: EP, MR, JC and ART. Funding acquisition: FA, BB, TM, VP, MY, JC and ART. Methodology: EP, MR, JC and ART. Project administration: JC and ART. Writing—original draft: EP, MR, JC and ART. Writing—review & editing: EP, MR, TM, BR, BB, FA, WJ, STE, TB and MY, JC and ART. All authors have read and approved the final version of the manuscript.

## Competing interests

TM has received personal fees from AMGEN, PIERRE FABRE, MSD, GALAPAGOS, SERVIER, MERCK SERONO and SANOFI, and TM declare participation on DSMB of the ongoing study PRODIGE 70 from FFCD, outside the submitted work. BR has received personal fees from AMGEN, ASTRAZENECA, BMS, CHUGAI, JANSSEN, LILLY, MSD, NOVARTIS, ROCHE, TAKEDA and declares no conflicts of interest in connection with this study. FA reports research grants or advisory boards paid to institutions from AstraZeneca, Daiichi Sankyo, Lilly, Novartis, BMS, Pfizer, Relay Tx and Roche and honorarium from Relay Therapeutics, Lilly, Astra Zeneca. WJ received personal fees from AstraZeneca, Daiichi Sankyo, Eisai, Novartis, Roche, Pfizer, Eli Lilly, MSD, BMS, Chugai, Seagen and Gilead. All other authors declare no conflicts of interest.

## Data and materials availability

The data that support these findings of the study are available upon request from the corresponding authors.

## Notes

### Competing Interest Statement

The authors have declared no competing interest.

