## Supplemental Figures for "Single-and double-strand circulating DNA fragmentomics for enhanced cancer detection performance"

### Slide 1
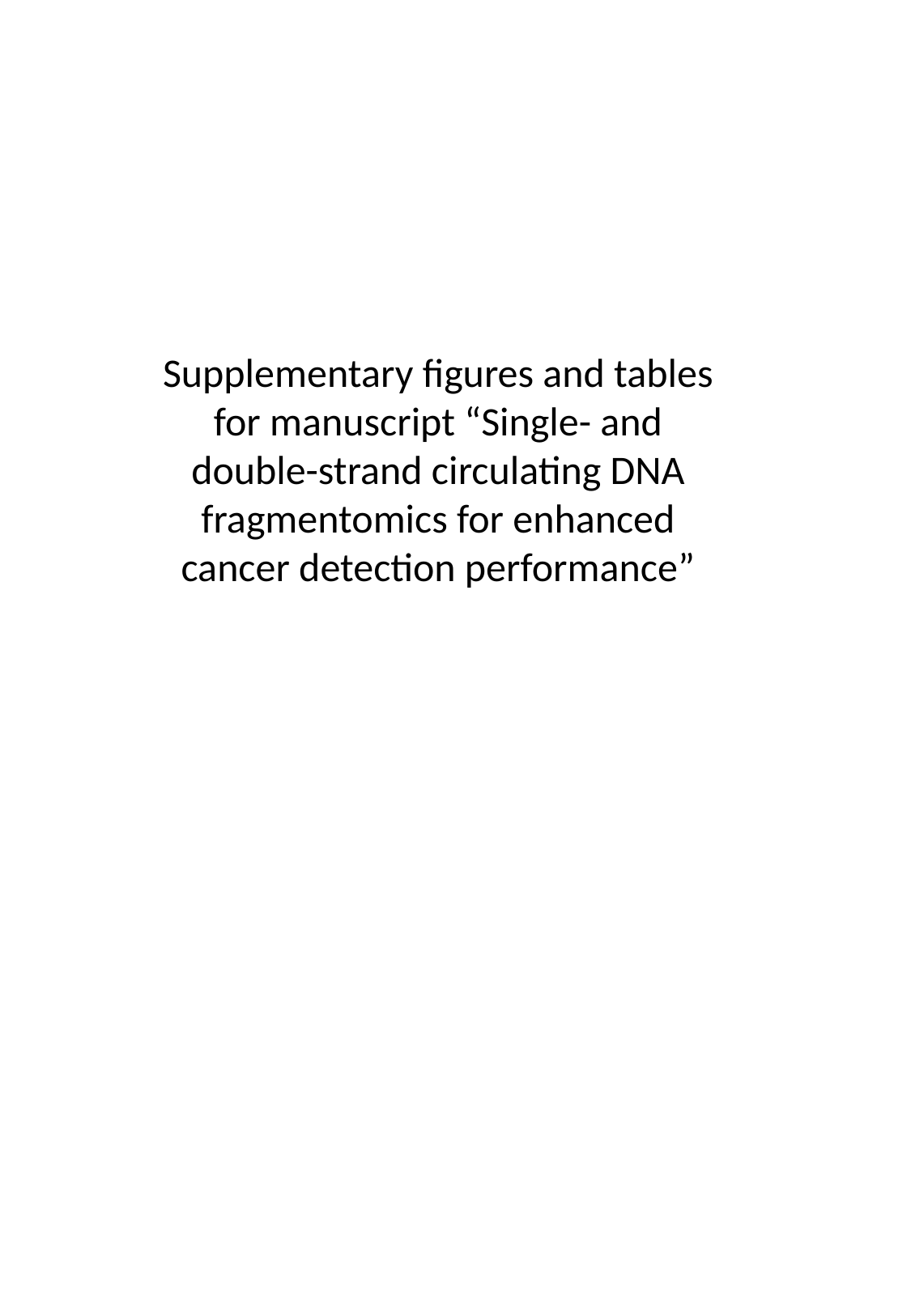

Supplementary figures and tables for manuscript “Single- and double-strand circulating DNA fragmentomics for enhanced cancer detection performance”

### Slide 2
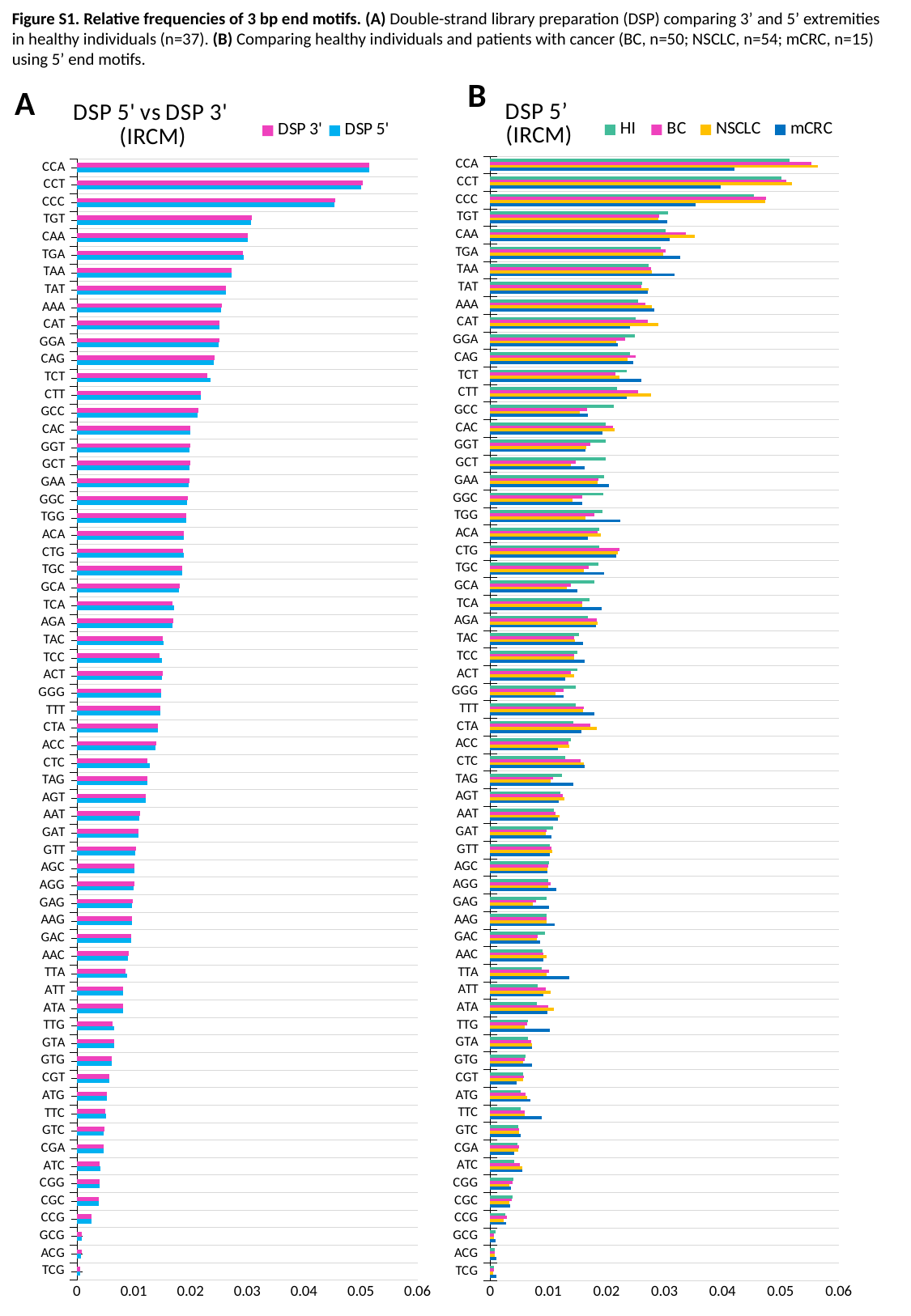

Figure S1. Relative frequencies of 3 bp end motifs. (A) Double-strand library preparation (DSP) comparing 3’ and 5’ extremities in healthy individuals (n=37). (B) Comparing healthy individuals and patients with cancer (BC, n=50; NSCLC, n=54; mCRC, n=15) using 5’ end motifs.
B
A
#### Chart: DSP 5' vs DSP 3'
(IRCM)
| Category | DSP 5' | DSP 3' |
|---|---|---|
| TCG | 0.000630913032183189 | 0.0006048581586794556 |
| ACG | 0.0007788186415914841 | 0.0007909093356480584 |
| GCG | 0.0008752247055238946 | 0.0008909745520540694 |
| CCG | 0.0025935839150228045 | 0.0026114674419079596 |
| CGC | 0.0038371033287111626 | 0.0038545586990866854 |
| CGG | 0.003961267612117466 | 0.00398099746889763 |
| ATC | 0.004129968528143292 | 0.004066238695227883 |
| CGA | 0.004703964639462332 | 0.004713343355727515 |
| GTC | 0.004763720763821107 | 0.004783919344970896 |
| TTC | 0.005172252541997134 | 0.004933936249823309 |
| ATG | 0.005217010881765665 | 0.0052760342946664195 |
| CGT | 0.005656012945472177 | 0.005751501912069717 |
| GTG | 0.006129383744612038 | 0.006176215255016045 |
| GTA | 0.0065408106410617005 | 0.006579422720365766 |
| TTG | 0.006552696333949614 | 0.006341356620072256 |
| ATA | 0.008081590801341493 | 0.008135432796264585 |
| ATT | 0.008145255324239464 | 0.008184520607455125 |
| TTA | 0.008810938151143014 | 0.008560904937356334 |
| AAC | 0.009044397156764817 | 0.00906473317166904 |
| GAC | 0.009486644914582317 | 0.009529214126390682 |
| AAG | 0.009694234355162314 | 0.00974578392233825 |
| GAG | 0.009737903903355122 | 0.009808498690688155 |
| AGG | 0.010043539725223974 | 0.010123434701736411 |
| AGC | 0.010095948685333513 | 0.010138151090775854 |
| GTT | 0.010259853584458814 | 0.010356790235279014 |
| GAT | 0.010805581075623323 | 0.010832566026507817 |
| AAT | 0.010925194348876472 | 0.01105692937587921 |
| AGT | 0.01208102348444286 | 0.012156133181663718 |
| TAG | 0.012383657445317704 | 0.012365753813832677 |
| CTC | 0.012851397645333931 | 0.012454262715794704 |
| ACC | 0.01384891452973676 | 0.013922301097788959 |
| CTA | 0.014307918872953409 | 0.014219938924672725 |
| TTT | 0.014757993539638423 | 0.014664199103957807 |
| GGG | 0.014781580595076409 | 0.01490048232926763 |
| ACT | 0.014958679266443332 | 0.01505386826031926 |
| TCC | 0.01503536732250551 | 0.0146097450408183 |
| TAC | 0.0152842148698834 | 0.015193926342820779 |
| AGA | 0.016801923445380222 | 0.016918783447681388 |
| TCA | 0.01707675663041124 | 0.016875534941169335 |
| GCA | 0.01798097167355866 | 0.01805223771932733 |
| TGC | 0.018597527936026315 | 0.018476620289247514 |
| CTG | 0.0188223200985531 | 0.018626400850072708 |
| ACA | 0.01882314992637162 | 0.01882896599090058 |
| TGG | 0.019330366008621994 | 0.019243569752283907 |
| GGC | 0.01943403716426345 | 0.019543761527920412 |
| GAA | 0.01966084364220491 | 0.01989227182538637 |
| GCT | 0.01985375806612627 | 0.01998118159448664 |
| GGT | 0.01985967711009642 | 0.01997587167753194 |
| CAC | 0.019940784815566468 | 0.019982184014159462 |
| GCC | 0.021252251095455726 | 0.021378279426890835 |
| CTT | 0.021882809068283632 | 0.021782170966953035 |
| TCT | 0.023469198008086684 | 0.022905299166274515 |
| CAG | 0.024096904197069113 | 0.02425829520353159 |
| GGA | 0.024964815804553484 | 0.025138755778379318 |
| CAT | 0.02504069403552931 | 0.02512139683296247 |
| AAA | 0.025416133298432668 | 0.025503114795515636 |
| TAT | 0.026183815127846666 | 0.026255633913392695 |
| TAA | 0.02721541891033746 | 0.027175581680104966 |
| TGA | 0.029414110668972152 | 0.029320995862483358 |
| CAA | 0.03017313428552358 | 0.030156065950988218 |
| TGT | 0.030678131759599468 | 0.03079497720496109 |
| CCC | 0.04540022232650994 | 0.045501616368423606 |
| CCT | 0.050119590160583226 | 0.05034683882807516 |
| CCA | 0.05154609288316478 | 0.051530289793405196 |
#### Chart: DSP 5’
(IRCM)
| Category | mCRC | NSCLC | BC | HI |
|---|---|---|---|---|
| TCG | 0.0011256999043050672 | 0.0005241251258677928 | 0.0005692127190800313 | 0.000630913032183189 |
| ACG | 0.000995429598058622 | 0.0007372345126597828 | 0.0007898587028200844 | 0.0007788186415914841 |
| GCG | 0.0009600702850958291 | 0.0005763178227160263 | 0.0007007999865431116 | 0.0008752247055238946 |
| CCG | 0.002749314878466809 | 0.002361758404446712 | 0.0028035062678605487 | 0.0025935839150228045 |
| CGC | 0.0033945989205330517 | 0.0033143272295148673 | 0.0037164831309052897 | 0.0038371033287111626 |
| CGG | 0.00358318174112614 | 0.003224789698778333 | 0.003864618480922581 | 0.003961267612117466 |
| ATC | 0.005577032918665712 | 0.005454781422571874 | 0.00506244563913846 | 0.004129968528143292 |
| CGA | 0.004180254624415456 | 0.004804149321332096 | 0.004963114241427675 | 0.004703964639462332 |
| GTC | 0.005260384660736397 | 0.004943394489104273 | 0.004905782042688778 | 0.004763720763821107 |
| TTC | 0.008903225401918896 | 0.005994121513735217 | 0.006007021827989726 | 0.005172252541997134 |
| ATG | 0.006900620527498098 | 0.0063204044241730135 | 0.006024779024878092 | 0.005217010881765665 |
| CGT | 0.00460039427576857 | 0.005679467379587055 | 0.005814009406823786 | 0.005656012945472177 |
| GTG | 0.007137691776154884 | 0.005689496134555481 | 0.005964798714418344 | 0.006129383744612038 |
| GTA | 0.007172205056879712 | 0.0072359179323360735 | 0.007116036194125098 | 0.0065408106410617005 |
| TTG | 0.010259129286687227 | 0.00591174516417606 | 0.006404752959701598 | 0.006552696333949614 |
| ATA | 0.009862281573933233 | 0.010933781043014004 | 0.009994471703102071 | 0.008081590801341493 |
| ATT | 0.009191994431369022 | 0.010455464575489527 | 0.009595942745957611 | 0.008145255324239464 |
| TTA | 0.013619553333906092 | 0.00973818447877102 | 0.010103972033586697 | 0.008810938151143014 |
| AAC | 0.009148024192972134 | 0.009644623227778803 | 0.009200774818280737 | 0.009044397156764817 |
| GAC | 0.008620715055085323 | 0.007992984062046569 | 0.008221575904227695 | 0.009486644914582317 |
| AAG | 0.011146501351790964 | 0.00976311507363386 | 0.009639474931900477 | 0.009694234355162314 |
| GAG | 0.010157698713995873 | 0.00733085353626098 | 0.00789616257204079 | 0.009737903903355122 |
| AGG | 0.01141040120907227 | 0.009923991508204685 | 0.0103696767738845 | 0.010043539725223974 |
| AGC | 0.00983101269507995 | 0.009796324135237181 | 0.009980574520015254 | 0.010095948685333513 |
| GTT | 0.01026529578802421 | 0.010702873464699376 | 0.010501249238989221 | 0.010259853584458814 |
| GAT | 0.01058420859168881 | 0.009611406255564062 | 0.009675632822609468 | 0.010805581075623323 |
| AAT | 0.011620074159697064 | 0.011906505556649027 | 0.011233168891440026 | 0.010925194348876472 |
| AGT | 0.011800074067085919 | 0.012761337837412031 | 0.012490225832687859 | 0.01208102348444286 |
| TAG | 0.01431186400450214 | 0.01035042412163242 | 0.010745636453597336 | 0.012383657445317704 |
| CTC | 0.016246002019670365 | 0.016057817384848412 | 0.015563851689765098 | 0.012851397645333931 |
| ACC | 0.011597197058503131 | 0.013603324270462304 | 0.01340268960578677 | 0.01384891452973676 |
| CTA | 0.01576392284130633 | 0.01839560254285997 | 0.017292497291923416 | 0.014307918872953409 |
| TTT | 0.01794403505093029 | 0.0159108937233809 | 0.01605484936850293 | 0.014757993539638423 |
| GGG | 0.01263864598509614 | 0.011193094263225475 | 0.012590872641630049 | 0.014781580595076409 |
| ACT | 0.012886224815185284 | 0.0144005465456697 | 0.013943616759347006 | 0.014958679266443332 |
| TCC | 0.01626781690214979 | 0.014402140833540857 | 0.014466136566695603 | 0.01503536732250551 |
| TAC | 0.015979194547977374 | 0.014618498789792102 | 0.014419385715055334 | 0.0152842148698834 |
| AGA | 0.018190737488535835 | 0.018549702696819378 | 0.018317941391574877 | 0.016801923445380222 |
| TCA | 0.01920907250404282 | 0.015806316658970955 | 0.015894860711988357 | 0.01707675663041124 |
| GCA | 0.014987658269353301 | 0.013223703776790849 | 0.01390841066484586 | 0.01798097167355866 |
| TGC | 0.01959285029447478 | 0.016139933720694086 | 0.017004857223410594 | 0.018597527936026315 |
| CTG | 0.0216971262888237 | 0.021937274793500165 | 0.02222552196113538 | 0.0188223200985531 |
| ACA | 0.01679566466314981 | 0.01898592021314139 | 0.01853151469668831 | 0.01882314992637162 |
| TGG | 0.022328799646573767 | 0.01636426534700691 | 0.017911217018741182 | 0.019330366008621994 |
| GGC | 0.01587407376636638 | 0.01418343061120555 | 0.015852450716277805 | 0.01943403716426345 |
| GAA | 0.0203836806425394 | 0.018523052771922538 | 0.0186965194878972 | 0.01966084364220491 |
| GCT | 0.016236613695790134 | 0.01394392889821972 | 0.014723197518947028 | 0.01985375806612627 |
| GGT | 0.01641176371790534 | 0.01650225816376825 | 0.017289094378540613 | 0.01985967711009642 |
| CAC | 0.019251519869822552 | 0.021421015024723412 | 0.02107898002689368 | 0.019940784815566468 |
| GCC | 0.016843846867180788 | 0.015382946425398027 | 0.01666746188384138 | 0.021252251095455726 |
| CTT | 0.02352390451446709 | 0.027626894442400258 | 0.025506885078753223 | 0.021882809068283632 |
| TCT | 0.025964960608284435 | 0.022233638387048733 | 0.02161906117149678 | 0.023469198008086684 |
| CAG | 0.0246826968584172 | 0.02360818782699219 | 0.02504596061123824 | 0.024096904197069113 |
| GGA | 0.02190691898905306 | 0.021719008527109254 | 0.023163158754345336 | 0.024964815804553484 |
| CAT | 0.02410093196310597 | 0.028928271921191025 | 0.02714956469561456 | 0.02504069403552931 |
| AAA | 0.028236153620380382 | 0.027837764431322125 | 0.026762714101422863 | 0.025416133298432668 |
| TAT | 0.027096312919638824 | 0.027329914165016338 | 0.02601801183234214 | 0.026183815127846666 |
| TAA | 0.03174699269200892 | 0.027763427658719413 | 0.027618644651601357 | 0.02721541891033746 |
| TGA | 0.03278295343899936 | 0.029836187237670838 | 0.030225103170164185 | 0.029414110668972152 |
| CAA | 0.030971616358383084 | 0.03519177227871898 | 0.03369594589073905 | 0.03017313428552358 |
| TGT | 0.03041782136363921 | 0.028986414249479203 | 0.02916414724667263 | 0.030678131759599468 |
| CCC | 0.035331354065567776 | 0.047419009533959645 | 0.047540162529762266 | 0.04540022232650994 |
| CCT | 0.03963797228323475 | 0.05191474243699449 | 0.050947615190467845 | 0.050119590160583226 |
| CCA | 0.04210403036492915 | 0.056375199995488345 | 0.05535133917424809 | 0.05154609288316478 |

### Slide 3
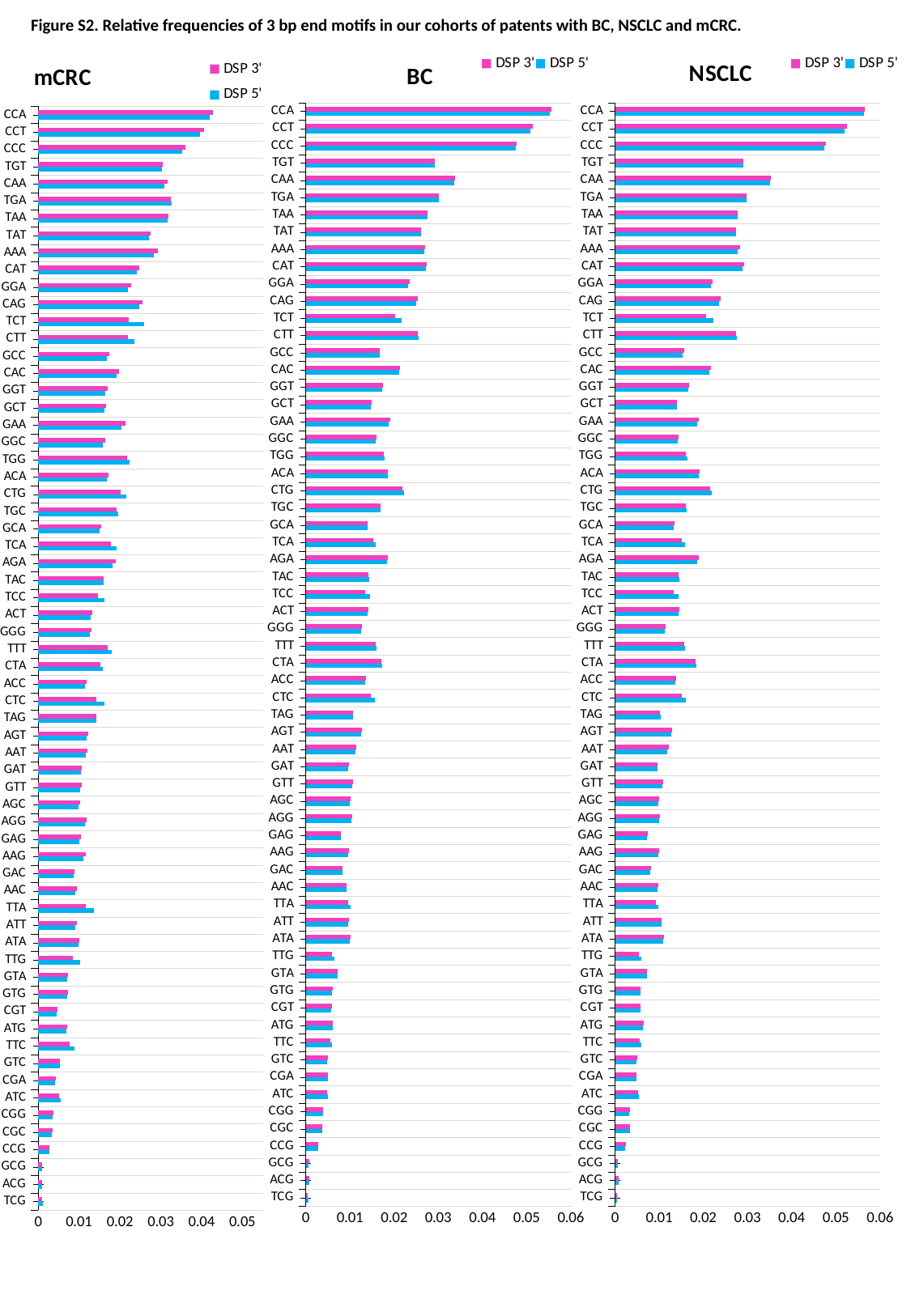

Figure S2. Relative frequencies of 3 bp end motifs in our cohorts of patents with BC, NSCLC and mCRC.
#### Chart: BC
| Category | DSP 5' | DSP 3' |
|---|---|---|
| TCG | 0.0005692127190800313 | 0.0004887070544483809 |
| ACG | 0.0007898587028200844 | 0.0008124065313506759 |
| GCG | 0.0007007999865431116 | 0.0007153974534691587 |
| CCG | 0.0028035062678605487 | 0.00284404567900817 |
| CGC | 0.0037164831309052897 | 0.003753874124723217 |
| CGG | 0.003864618480922581 | 0.003931233579796172 |
| ATC | 0.00506244563913846 | 0.0049101899187016405 |
| CGA | 0.004963114241427675 | 0.0049727914423337025 |
| GTC | 0.004905782042688778 | 0.004945900811557798 |
| TTC | 0.006007021827989726 | 0.00561168927791619 |
| ATG | 0.006024779024878092 | 0.006152863753376699 |
| CGT | 0.005814009406823786 | 0.005912924097146428 |
| GTG | 0.005964798714418344 | 0.006062160837566166 |
| GTA | 0.007116036194125098 | 0.007197549515421034 |
| TTG | 0.006404752959701598 | 0.005924886427451912 |
| ATA | 0.009994471703102071 | 0.010160155837259823 |
| ATT | 0.009595942745957611 | 0.00969052081178602 |
| TTA | 0.010103972033586697 | 0.009613200783093545 |
| AAC | 0.009200774818280737 | 0.009292789089661017 |
| GAC | 0.008221575904227695 | 0.008319928486433217 |
| AAG | 0.009639474931900477 | 0.009768018337156244 |
| GAG | 0.00789616257204079 | 0.008010236214241405 |
| AGG | 0.0103696767738845 | 0.01055169903471362 |
| AGC | 0.009980574520015254 | 0.010118559474041243 |
| GTT | 0.010501249238989221 | 0.010677911629783085 |
| GAT | 0.009675632822609468 | 0.00973578627891063 |
| AAT | 0.011233168891440026 | 0.01146059212783563 |
| AGT | 0.012490225832687859 | 0.012654773081289775 |
| TAG | 0.010745636453597336 | 0.010612664749938132 |
| CTC | 0.015563851689765098 | 0.014652610493030974 |
| ACC | 0.01340268960578677 | 0.013547941411730562 |
| CTA | 0.017292497291923416 | 0.01715606356685039 |
| TTT | 0.01605484936850293 | 0.015828932838847717 |
| GGG | 0.012590872641630049 | 0.012773622070209156 |
| ACT | 0.013943616759347006 | 0.01412079156003817 |
| TCC | 0.014466136566695603 | 0.01342988486964057 |
| TAC | 0.014419385715055334 | 0.014265508175668552 |
| AGA | 0.018317941391574877 | 0.018574071422256713 |
| TCA | 0.015894860711988357 | 0.015286937605676237 |
| GCA | 0.01390841066484586 | 0.014054313412472654 |
| TGC | 0.017004857223410594 | 0.01684530407421803 |
| CTG | 0.02222552196113538 | 0.021893019831998775 |
| ACA | 0.01853151469668831 | 0.018667113453148206 |
| TGG | 0.017911217018741182 | 0.017715671118609153 |
| GGC | 0.015852450716277805 | 0.016029898800168942 |
| GAA | 0.0186965194878972 | 0.019043871525329395 |
| GCT | 0.014723197518947028 | 0.014901803971056272 |
| GGT | 0.017289094378540613 | 0.01750694261907877 |
| CAC | 0.02107898002689368 | 0.021239902309101225 |
| GCC | 0.01666746188384138 | 0.01683943708115261 |
| CTT | 0.025506885078753223 | 0.02534129855794645 |
| TCT | 0.02161906117149678 | 0.020298272816544203 |
| CAG | 0.02504596061123824 | 0.025392398389443384 |
| GGA | 0.023163158754345336 | 0.023458734500928502 |
| CAT | 0.02714956469561456 | 0.027397304136414752 |
| AAA | 0.026762714101422863 | 0.027085098202422008 |
| TAT | 0.02601801183234214 | 0.02608246855642584 |
| TAA | 0.027618644651601357 | 0.027560516648546023 |
| TGA | 0.030225103170164185 | 0.030156052840920555 |
| CAA | 0.03369594589073905 | 0.03381503580937266 |
| TGT | 0.02916414724667263 | 0.029305926615454402 |
| CCC | 0.047540162529762266 | 0.0478145499479427 |
| CCT | 0.050947615190467845 | 0.05145049125119308 |
| CCA | 0.05535133917424809 | 0.055564753075751615 |
#### Chart: NSCLC
| Category | DSP 5' | DSP 3' |
|---|---|---|
| TCG | 0.0005241251258677928 | 0.0004275425439704377 |
| ACG | 0.0007372345126597828 | 0.0007521248650899966 |
| GCG | 0.0005763178227160263 | 0.0005934701519974724 |
| CCG | 0.002361758404446712 | 0.002401100530998106 |
| CGC | 0.0033143272295148673 | 0.0033495970884597657 |
| CGG | 0.003224789698778333 | 0.0032956127438644083 |
| ATC | 0.005454781422571874 | 0.005275476284208415 |
| CGA | 0.004804149321332096 | 0.0048283251468061025 |
| GTC | 0.004943394489104273 | 0.005022966344904971 |
| TTC | 0.005994121513735217 | 0.00554277085713259 |
| ATG | 0.0063204044241730135 | 0.006474048419616525 |
| CGT | 0.005679467379587055 | 0.005805588732294914 |
| GTG | 0.005689496134555481 | 0.005805105781883907 |
| GTA | 0.0072359179323360735 | 0.007330168436108403 |
| TTG | 0.00591174516417606 | 0.005334424294458536 |
| ATA | 0.010933781043014004 | 0.011078243763713164 |
| ATT | 0.010455464575489527 | 0.01058095621830667 |
| TTA | 0.00973818447877102 | 0.009167760557761782 |
| AAC | 0.009644623227778803 | 0.009777439553482955 |
| GAC | 0.007992984062046569 | 0.008097970321704707 |
| AAG | 0.00976311507363386 | 0.009923824276985443 |
| GAG | 0.00733085353626098 | 0.007464319397491756 |
| AGG | 0.009923991508204685 | 0.01011956315451662 |
| AGC | 0.009796324135237181 | 0.009951190606311165 |
| GTT | 0.010702873464699376 | 0.010890143714710162 |
| GAT | 0.009611406255564062 | 0.009683455647289068 |
| AAT | 0.011906505556649027 | 0.012167934475059386 |
| AGT | 0.012761337837412031 | 0.013005435508276314 |
| TAG | 0.01035042412163242 | 0.010188139050073418 |
| CTC | 0.016057817384848412 | 0.01506474639650307 |
| ACC | 0.013603324270462304 | 0.013772989147997171 |
| CTA | 0.01839560254285997 | 0.01827491139340273 |
| TTT | 0.0159108937233809 | 0.015635629415084024 |
| GGG | 0.011193094263225475 | 0.01136346842283191 |
| ACT | 0.0144005465456697 | 0.01460150166526567 |
| TCC | 0.014402140833540857 | 0.013204597355810388 |
| TAC | 0.014618498789792102 | 0.014468368047282831 |
| AGA | 0.018549702696819378 | 0.018903841980919656 |
| TCA | 0.015806316658970955 | 0.015086428299146816 |
| GCA | 0.013223703776790849 | 0.013384527077021964 |
| TGC | 0.016139933720694086 | 0.015961895885040767 |
| CTG | 0.021937274793500165 | 0.021619923624682364 |
| ACA | 0.01898592021314139 | 0.01912970677737087 |
| TGG | 0.01636426534700691 | 0.016105563703758714 |
| GGC | 0.01418343061120555 | 0.014378491033715714 |
| GAA | 0.018523052771922538 | 0.0189224905590997 |
| GCT | 0.01394392889821972 | 0.014108440040675158 |
| GGT | 0.01650225816376825 | 0.01673015141517147 |
| CAC | 0.021421015024723412 | 0.021634627210078135 |
| GCC | 0.015382946425398027 | 0.015586464301665912 |
| CTT | 0.027626894442400258 | 0.027435524686483432 |
| TCT | 0.022233638387048733 | 0.020609471613712245 |
| CAG | 0.02360818782699219 | 0.02394728849973935 |
| GGA | 0.021719008527109254 | 0.022057903947601543 |
| CAT | 0.028928271921191025 | 0.029199052391079405 |
| AAA | 0.027837764431322125 | 0.028271365067740303 |
| TAT | 0.027329914165016338 | 0.02738793437133286 |
| TAA | 0.027763427658719413 | 0.027692151745769278 |
| TGA | 0.029836187237670838 | 0.02979687204573852 |
| CAA | 0.03519177227871898 | 0.03536645529973982 |
| TGT | 0.028986414249479203 | 0.02910400775789784 |
| CCC | 0.047419009533959645 | 0.04771850500542298 |
| CCT | 0.05191474243699449 | 0.05252037466185259 |
| CCA | 0.056375199995488345 | 0.05661963068588761 |
#### Chart: mCRC
| Category | DSP 5' | DSP 3' |
|---|---|---|
| TCG | 0.0011256999043050672 | 0.0009030471878754381 |
| ACG | 0.000995429598058622 | 0.0010485704134020827 |
| GCG | 0.0009600702850958291 | 0.0010120345977483675 |
| CCG | 0.002749314878466809 | 0.0028348756847951517 |
| CGC | 0.0033945989205330517 | 0.00352408848902796 |
| CGG | 0.00358318174112614 | 0.0037566126450318603 |
| ATC | 0.005577032918665712 | 0.005156220261037792 |
| CGA | 0.004180254624415456 | 0.0042891224316772185 |
| GTC | 0.005260384660736397 | 0.005424693610546885 |
| TTC | 0.008903225401918896 | 0.007782908007381382 |
| ATG | 0.006900620527498098 | 0.007185842277229788 |
| CGT | 0.00460039427576857 | 0.004798650605309302 |
| GTG | 0.007137691776154884 | 0.0074127512241840155 |
| GTA | 0.007172205056879712 | 0.007387953298811404 |
| TTG | 0.010259129286687227 | 0.008461534116533408 |
| ATA | 0.009862281573933233 | 0.010161000309788832 |
| ATT | 0.009191994431369022 | 0.009432476960201368 |
| TTA | 0.013619553333906092 | 0.011685708914029976 |
| AAC | 0.009148024192972134 | 0.009489153439871854 |
| GAC | 0.008620715055085323 | 0.008931754704512691 |
| AAG | 0.011146501351790964 | 0.0116694050998053 |
| GAG | 0.010157698713995873 | 0.01058163388348936 |
| AGG | 0.01141040120907227 | 0.011904624226813513 |
| AGC | 0.00983101269507995 | 0.010268924522822425 |
| GTT | 0.01026529578802421 | 0.010644366972000146 |
| GAT | 0.01058420859168881 | 0.010747154951302287 |
| AAT | 0.011620074159697064 | 0.012158206553537729 |
| AGT | 0.011800074067085919 | 0.012275336630291652 |
| TAG | 0.01431186400450214 | 0.014188258918778366 |
| CTC | 0.016246002019670365 | 0.014179817497958488 |
| ACC | 0.011597197058503131 | 0.011917927904461167 |
| CTA | 0.01576392284130633 | 0.015290805908680153 |
| TTT | 0.01794403505093029 | 0.01702719204235067 |
| GGG | 0.01263864598509614 | 0.01307230345521417 |
| ACT | 0.012886224815185284 | 0.013240830468897567 |
| TCC | 0.01626781690214979 | 0.014700800506570218 |
| TAC | 0.015979194547977374 | 0.015990154401810786 |
| AGA | 0.018190737488535835 | 0.018949088700234172 |
| TCA | 0.01920907250404282 | 0.017813960050888863 |
| GCA | 0.014987658269353301 | 0.015418925022039998 |
| TGC | 0.01959285029447478 | 0.019203247587010055 |
| CTG | 0.0216971262888237 | 0.020233714428894736 |
| ACA | 0.01679566466314981 | 0.017198528024006607 |
| TGG | 0.022328799646573767 | 0.021804099265060158 |
| GGC | 0.01587407376636638 | 0.016405277801483497 |
| GAA | 0.0203836806425394 | 0.021344918874365786 |
| GCT | 0.016236613695790134 | 0.01670202605040156 |
| GGT | 0.01641176371790534 | 0.01696347638023962 |
| CAC | 0.019251519869822552 | 0.019859374345935536 |
| GCC | 0.016843846867180788 | 0.01736976195705723 |
| CTT | 0.02352390451446709 | 0.022015959002649483 |
| TCT | 0.025964960608284435 | 0.022231973187703168 |
| CAG | 0.0246826968584172 | 0.025571709986402246 |
| GGA | 0.02190691898905306 | 0.022704572922662664 |
| CAT | 0.02410093196310597 | 0.024783032240433874 |
| AAA | 0.028236153620380382 | 0.02940434576480173 |
| TAT | 0.027096312919638824 | 0.027468776795560243 |
| TAA | 0.03174699269200892 | 0.03182307874452346 |
| TGA | 0.03278295343899936 | 0.03254067191417652 |
| CAA | 0.030971616358383084 | 0.03176047508708167 |
| TGT | 0.03041782136363921 | 0.030574066304397234 |
| CCC | 0.035331354065567776 | 0.03597184689751529 |
| CCT | 0.03963797228323475 | 0.04053332596867348 |
| CCA | 0.04210403036492915 | 0.04281302357202034 |

### Slide 4
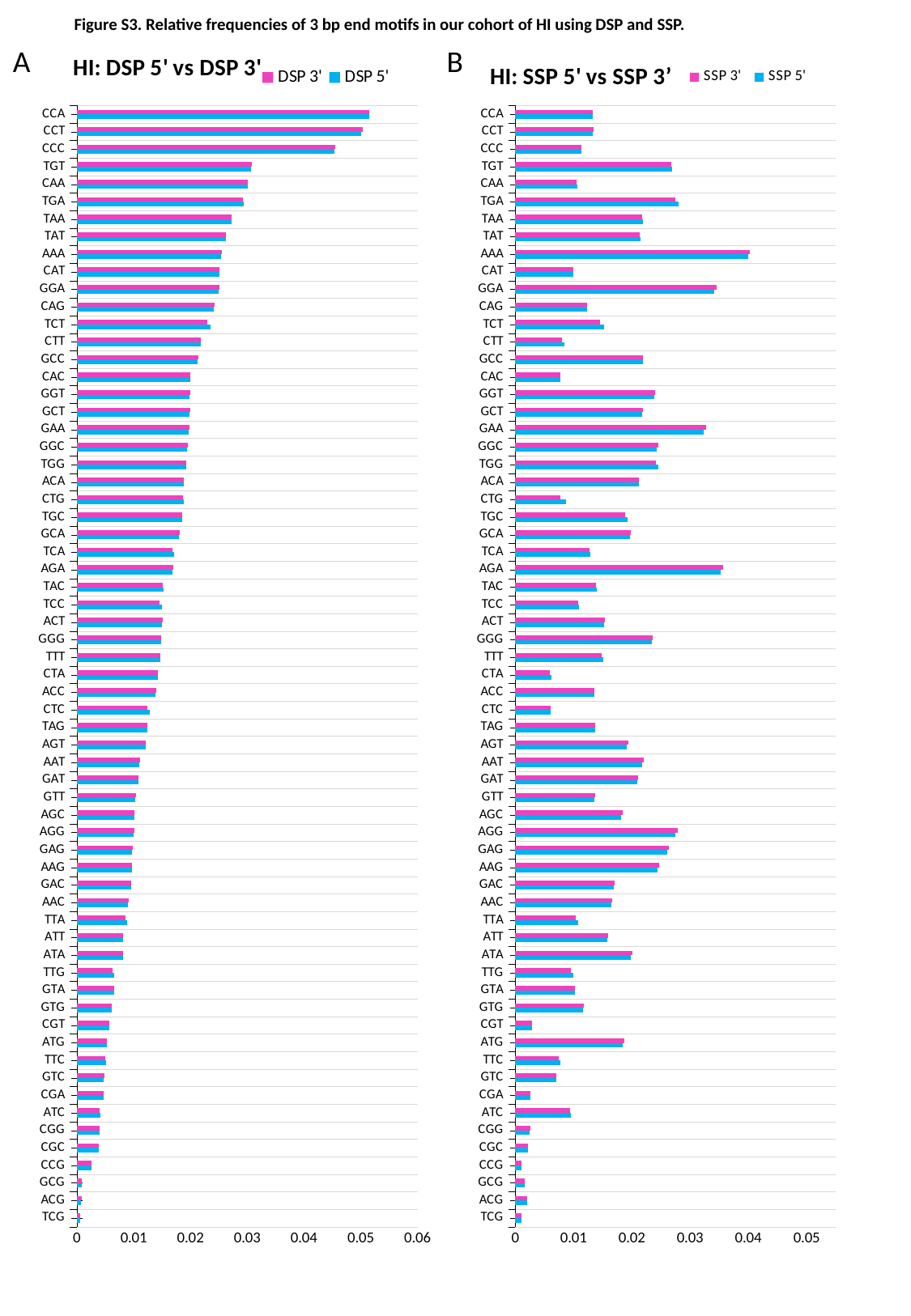

Figure S3. Relative frequencies of 3 bp end motifs in our cohort of HI using DSP and SSP.
A
B
#### Chart: HI: DSP 5' vs DSP 3'
| Category | DSP 5' | DSP 3' |
|---|---|---|
| TCG | 0.000630913032183189 | 0.0006048581586794556 |
| ACG | 0.0007788186415914841 | 0.0007909093356480584 |
| GCG | 0.0008752247055238946 | 0.0008909745520540694 |
| CCG | 0.0025935839150228045 | 0.0026114674419079596 |
| CGC | 0.0038371033287111626 | 0.0038545586990866854 |
| CGG | 0.003961267612117466 | 0.00398099746889763 |
| ATC | 0.004129968528143292 | 0.004066238695227883 |
| CGA | 0.004703964639462332 | 0.004713343355727515 |
| GTC | 0.004763720763821107 | 0.004783919344970896 |
| TTC | 0.005172252541997134 | 0.004933936249823309 |
| ATG | 0.005217010881765665 | 0.0052760342946664195 |
| CGT | 0.005656012945472177 | 0.005751501912069717 |
| GTG | 0.006129383744612038 | 0.006176215255016045 |
| GTA | 0.0065408106410617005 | 0.006579422720365766 |
| TTG | 0.006552696333949614 | 0.006341356620072256 |
| ATA | 0.008081590801341493 | 0.008135432796264585 |
| ATT | 0.008145255324239464 | 0.008184520607455125 |
| TTA | 0.008810938151143014 | 0.008560904937356334 |
| AAC | 0.009044397156764817 | 0.00906473317166904 |
| GAC | 0.009486644914582317 | 0.009529214126390682 |
| AAG | 0.009694234355162314 | 0.00974578392233825 |
| GAG | 0.009737903903355122 | 0.009808498690688155 |
| AGG | 0.010043539725223974 | 0.010123434701736411 |
| AGC | 0.010095948685333513 | 0.010138151090775854 |
| GTT | 0.010259853584458814 | 0.010356790235279014 |
| GAT | 0.010805581075623323 | 0.010832566026507817 |
| AAT | 0.010925194348876472 | 0.01105692937587921 |
| AGT | 0.01208102348444286 | 0.012156133181663718 |
| TAG | 0.012383657445317704 | 0.012365753813832677 |
| CTC | 0.012851397645333931 | 0.012454262715794704 |
| ACC | 0.01384891452973676 | 0.013922301097788959 |
| CTA | 0.014307918872953409 | 0.014219938924672725 |
| TTT | 0.014757993539638423 | 0.014664199103957807 |
| GGG | 0.014781580595076409 | 0.01490048232926763 |
| ACT | 0.014958679266443332 | 0.01505386826031926 |
| TCC | 0.01503536732250551 | 0.0146097450408183 |
| TAC | 0.0152842148698834 | 0.015193926342820779 |
| AGA | 0.016801923445380222 | 0.016918783447681388 |
| TCA | 0.01707675663041124 | 0.016875534941169335 |
| GCA | 0.01798097167355866 | 0.01805223771932733 |
| TGC | 0.018597527936026315 | 0.018476620289247514 |
| CTG | 0.0188223200985531 | 0.018626400850072708 |
| ACA | 0.01882314992637162 | 0.01882896599090058 |
| TGG | 0.019330366008621994 | 0.019243569752283907 |
| GGC | 0.01943403716426345 | 0.019543761527920412 |
| GAA | 0.01966084364220491 | 0.01989227182538637 |
| GCT | 0.01985375806612627 | 0.01998118159448664 |
| GGT | 0.01985967711009642 | 0.01997587167753194 |
| CAC | 0.019940784815566468 | 0.019982184014159462 |
| GCC | 0.021252251095455726 | 0.021378279426890835 |
| CTT | 0.021882809068283632 | 0.021782170966953035 |
| TCT | 0.023469198008086684 | 0.022905299166274515 |
| CAG | 0.024096904197069113 | 0.02425829520353159 |
| GGA | 0.024964815804553484 | 0.025138755778379318 |
| CAT | 0.02504069403552931 | 0.02512139683296247 |
| AAA | 0.025416133298432668 | 0.025503114795515636 |
| TAT | 0.026183815127846666 | 0.026255633913392695 |
| TAA | 0.02721541891033746 | 0.027175581680104966 |
| TGA | 0.029414110668972152 | 0.029320995862483358 |
| CAA | 0.03017313428552358 | 0.030156065950988218 |
| TGT | 0.030678131759599468 | 0.03079497720496109 |
| CCC | 0.04540022232650994 | 0.045501616368423606 |
| CCT | 0.050119590160583226 | 0.05034683882807516 |
| CCA | 0.05154609288316478 | 0.051530289793405196 |
#### Chart: HI: SSP 5' vs SSP 3’
| Category | SSP 5' | SSP 3' |
|---|---|---|
| TCG | 0.001031759903875992 | 0.00100940080521349 |
| ACG | 0.0020516373053169707 | 0.002081717441789616 |
| GCG | 0.0016216281133382042 | 0.0016444286612152463 |
| CCG | 0.0010161333791887746 | 0.0010172429067202046 |
| CGC | 0.002167801992980307 | 0.0021695088957972715 |
| CGG | 0.0025133927218297376 | 0.0025343410487256844 |
| ATC | 0.00950727860860273 | 0.00946389741119394 |
| CGA | 0.0025532236105255427 | 0.0025569834349536565 |
| GTC | 0.006986182895734692 | 0.007033465376727488 |
| TTC | 0.007677749792470413 | 0.007414470333322514 |
| ATG | 0.01843266054071008 | 0.01870453507748164 |
| CGT | 0.002799958677236697 | 0.0028227704544536323 |
| GTG | 0.011677037771451158 | 0.011744780683584559 |
| GTA | 0.010168751758189256 | 0.010230936775328305 |
| TTG | 0.009926691008239468 | 0.009547149272667444 |
| ATA | 0.01986552828433844 | 0.020101831720630257 |
| ATT | 0.015861078197914536 | 0.015991836893671337 |
| TTA | 0.01084749096823796 | 0.01035386441670134 |
| AAC | 0.01655445131433323 | 0.016690199698662946 |
| GAC | 0.01684324949849817 | 0.016979101752257163 |
| AAG | 0.024475266376885893 | 0.024723124665688177 |
| GAG | 0.02602681223722741 | 0.026289071514506115 |
| AGG | 0.027498741648158073 | 0.02785764605618124 |
| AGC | 0.018219213681852538 | 0.01840196392999955 |
| GTT | 0.01355429060761149 | 0.013696904879583093 |
| GAT | 0.020991104573245503 | 0.0210531043218857 |
| AAT | 0.021728707498367975 | 0.0220890748893402 |
| AGT | 0.019188897351632195 | 0.019405402697152418 |
| TAG | 0.013773095289780744 | 0.01368896657534788 |
| CTC | 0.006133220025675457 | 0.005999831331330767 |
| ACC | 0.013525208920583875 | 0.013633494268421826 |
| CTA | 0.006143402113603013 | 0.005877612039590751 |
| TTT | 0.015094865937565644 | 0.014785347216594063 |
| GGG | 0.02337011071593184 | 0.023551147850291786 |
| ACT | 0.01526276165420827 | 0.01539191716000067 |
| TCC | 0.010970762615616716 | 0.01082616168462408 |
| TAC | 0.013927738347541178 | 0.013817519912225218 |
| AGA | 0.03529025165484158 | 0.03572831621660438 |
| TCA | 0.012802459792491186 | 0.01275240770377763 |
| GCA | 0.01965242188544361 | 0.019773591227677367 |
| TGC | 0.01919569903165412 | 0.018795184716141798 |
| CTG | 0.008699062912093684 | 0.007752396663298388 |
| ACA | 0.021175132538896753 | 0.02127724586193716 |
| TGG | 0.024523003029700865 | 0.024085583910538194 |
| GGC | 0.024325384095141116 | 0.02449468459863585 |
| GAA | 0.03231873162492096 | 0.032819441118613187 |
| GCT | 0.021728440329786755 | 0.02185247895114298 |
| GGT | 0.023815838463226418 | 0.024012621933397163 |
| CAC | 0.007689080438859095 | 0.007718767193074706 |
| GCC | 0.021850960266621616 | 0.021969436261173248 |
| CTT | 0.008405926991310006 | 0.008039652447913768 |
| TCT | 0.015250541767486947 | 0.014581805186592476 |
| CAG | 0.01225875696897176 | 0.012324202051662509 |
| GGA | 0.03420425393980899 | 0.03459750400835909 |
| CAT | 0.009925209600265694 | 0.009912152116273484 |
| AAA | 0.03999614191115867 | 0.040318575478691734 |
| TAT | 0.02148433646387837 | 0.02141616596543629 |
| TAA | 0.021962812173272835 | 0.021714731416867278 |
| TGA | 0.027974451463310963 | 0.027502943272622017 |
| CAA | 0.010611616463259019 | 0.010527302921504191 |
| TGT | 0.026859885772914396 | 0.026731775773542635 |
| CCC | 0.011311893822613378 | 0.011379933296599158 |
| CCT | 0.013341200861014545 | 0.01339129998585879 |
| CCA | 0.013358619798556466 | 0.01334904566820323 |

### Slide 5
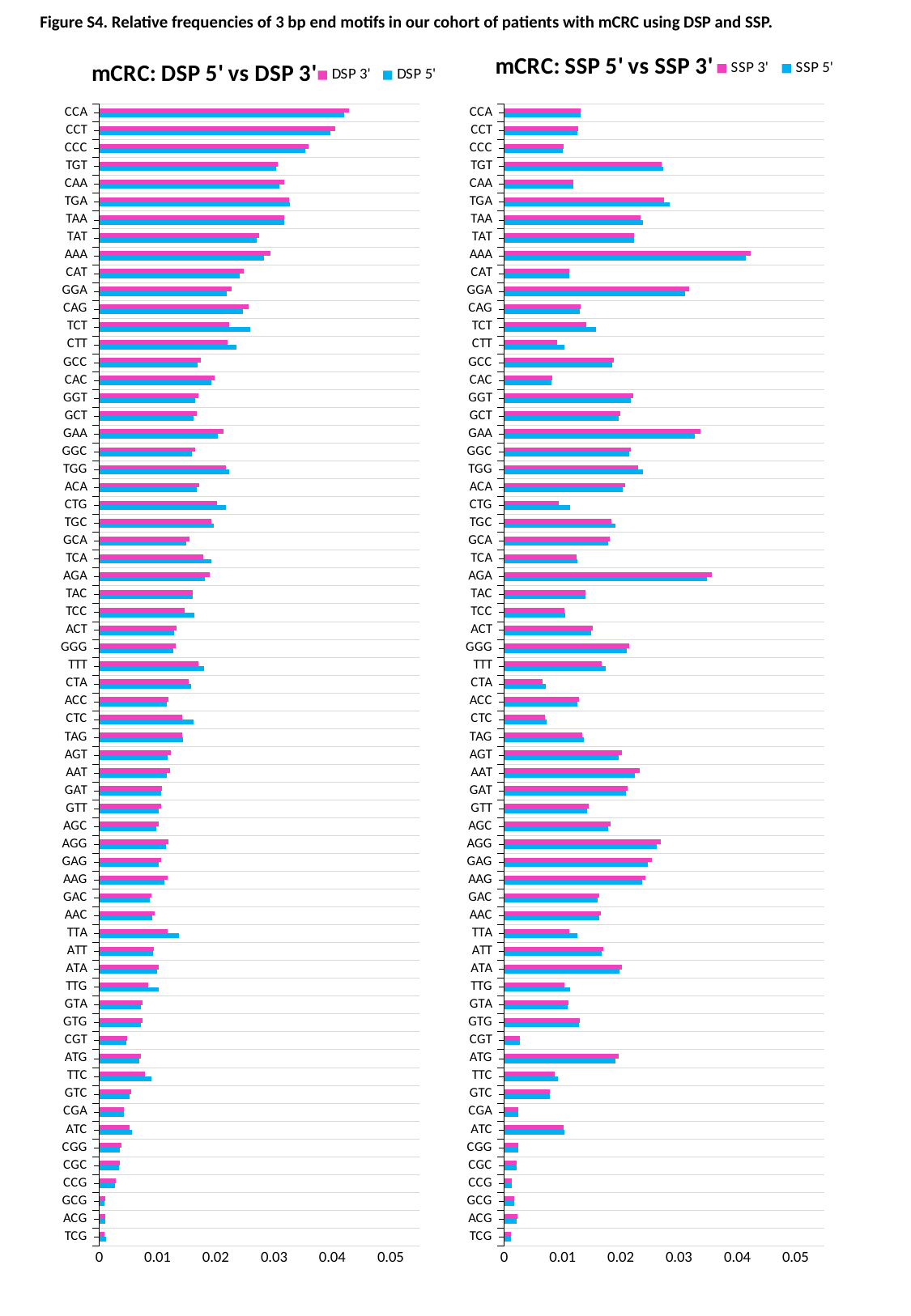

Figure S4. Relative frequencies of 3 bp end motifs in our cohort of patients with mCRC using DSP and SSP.
#### Chart: mCRC: DSP 5' vs DSP 3'
| Category | DSP 5' | DSP 3' |
|---|---|---|
| TCG | 0.0011256999043050672 | 0.0009030471878754381 |
| ACG | 0.000995429598058622 | 0.0010485704134020827 |
| GCG | 0.0009600702850958291 | 0.0010120345977483675 |
| CCG | 0.002749314878466809 | 0.0028348756847951517 |
| CGC | 0.0033945989205330517 | 0.00352408848902796 |
| CGG | 0.00358318174112614 | 0.0037566126450318603 |
| ATC | 0.005577032918665712 | 0.005156220261037792 |
| CGA | 0.004180254624415456 | 0.0042891224316772185 |
| GTC | 0.005260384660736397 | 0.005424693610546885 |
| TTC | 0.008903225401918896 | 0.007782908007381382 |
| ATG | 0.006900620527498098 | 0.007185842277229788 |
| CGT | 0.00460039427576857 | 0.004798650605309302 |
| GTG | 0.007137691776154884 | 0.0074127512241840155 |
| GTA | 0.007172205056879712 | 0.007387953298811404 |
| TTG | 0.010259129286687227 | 0.008461534116533408 |
| ATA | 0.009862281573933233 | 0.010161000309788832 |
| ATT | 0.009191994431369022 | 0.009432476960201368 |
| TTA | 0.013619553333906092 | 0.011685708914029976 |
| AAC | 0.009148024192972134 | 0.009489153439871854 |
| GAC | 0.008620715055085323 | 0.008931754704512691 |
| AAG | 0.011146501351790964 | 0.0116694050998053 |
| GAG | 0.010157698713995873 | 0.01058163388348936 |
| AGG | 0.01141040120907227 | 0.011904624226813513 |
| AGC | 0.00983101269507995 | 0.010268924522822425 |
| GTT | 0.01026529578802421 | 0.010644366972000146 |
| GAT | 0.01058420859168881 | 0.010747154951302287 |
| AAT | 0.011620074159697064 | 0.012158206553537729 |
| AGT | 0.011800074067085919 | 0.012275336630291652 |
| TAG | 0.01431186400450214 | 0.014188258918778366 |
| CTC | 0.016246002019670365 | 0.014179817497958488 |
| ACC | 0.011597197058503131 | 0.011917927904461167 |
| CTA | 0.01576392284130633 | 0.015290805908680153 |
| TTT | 0.01794403505093029 | 0.01702719204235067 |
| GGG | 0.01263864598509614 | 0.01307230345521417 |
| ACT | 0.012886224815185284 | 0.013240830468897567 |
| TCC | 0.01626781690214979 | 0.014700800506570218 |
| TAC | 0.015979194547977374 | 0.015990154401810786 |
| AGA | 0.018190737488535835 | 0.018949088700234172 |
| TCA | 0.01920907250404282 | 0.017813960050888863 |
| GCA | 0.014987658269353301 | 0.015418925022039998 |
| TGC | 0.01959285029447478 | 0.019203247587010055 |
| CTG | 0.0216971262888237 | 0.020233714428894736 |
| ACA | 0.01679566466314981 | 0.017198528024006607 |
| TGG | 0.022328799646573767 | 0.021804099265060158 |
| GGC | 0.01587407376636638 | 0.016405277801483497 |
| GAA | 0.0203836806425394 | 0.021344918874365786 |
| GCT | 0.016236613695790134 | 0.01670202605040156 |
| GGT | 0.01641176371790534 | 0.01696347638023962 |
| CAC | 0.019251519869822552 | 0.019859374345935536 |
| GCC | 0.016843846867180788 | 0.01736976195705723 |
| CTT | 0.02352390451446709 | 0.022015959002649483 |
| TCT | 0.025964960608284435 | 0.022231973187703168 |
| CAG | 0.0246826968584172 | 0.025571709986402246 |
| GGA | 0.02190691898905306 | 0.022704572922662664 |
| CAT | 0.02410093196310597 | 0.024783032240433874 |
| AAA | 0.028236153620380382 | 0.02940434576480173 |
| TAT | 0.027096312919638824 | 0.027468776795560243 |
| TAA | 0.03174699269200892 | 0.03182307874452346 |
| TGA | 0.03278295343899936 | 0.03254067191417652 |
| CAA | 0.030971616358383084 | 0.03176047508708167 |
| TGT | 0.03041782136363921 | 0.030574066304397234 |
| CCC | 0.035331354065567776 | 0.03597184689751529 |
| CCT | 0.03963797228323475 | 0.04053332596867348 |
| CCA | 0.04210403036492915 | 0.04281302357202034 |
#### Chart: mCRC: SSP 5' vs SSP 3'
| Category | SSP 5' | SSP 3' |
|---|---|---|
| TCG | 0.0011708624653753385 | 0.0010945887920592442 |
| ACG | 0.0021856709012487057 | 0.0022409066946741806 |
| GCG | 0.0017024410159432353 | 0.0017301651873302423 |
| CCG | 0.0012380341511327956 | 0.0012450348339000548 |
| CGC | 0.002068894827934708 | 0.0020934409700273738 |
| CGG | 0.002416754964140893 | 0.0024692983644430864 |
| ATC | 0.010287571984399077 | 0.010182415284715455 |
| CGA | 0.0024463744084479316 | 0.002453486666927766 |
| GTC | 0.007795995651756647 | 0.007884549167599864 |
| TTC | 0.009255763615823486 | 0.008686870443379366 |
| ATG | 0.019036437927387354 | 0.019609161356254456 |
| CGT | 0.0026298734307367896 | 0.002679602335170819 |
| GTG | 0.012798469556950501 | 0.01294359326574286 |
| GTA | 0.01093070833168127 | 0.011053985659588435 |
| TTG | 0.011365899990486436 | 0.010279300671277603 |
| ATA | 0.019736078962638282 | 0.020151690089787774 |
| ATT | 0.016760085465544826 | 0.01705586539901172 |
| TTA | 0.012566042640707481 | 0.011143314596004946 |
| AAC | 0.016256839759456725 | 0.016605599627582537 |
| GAC | 0.01603857411795272 | 0.01634203768160577 |
| AAG | 0.023648054541405308 | 0.02425820816814285 |
| GAG | 0.024686080296445995 | 0.025276570450914707 |
| AGG | 0.026096938423310394 | 0.02684816210871009 |
| AGC | 0.017864108750969835 | 0.01830991011553874 |
| GTT | 0.014269274366584586 | 0.014523900987058793 |
| GAT | 0.020880718831179736 | 0.02118375292010348 |
| AAT | 0.0224815813775653 | 0.023273517221464073 |
| AGT | 0.019633691482187472 | 0.02016488637460025 |
| TAG | 0.01366233629103045 | 0.013406169741426315 |
| CTC | 0.007217973897222636 | 0.006939955711222017 |
| ACC | 0.01253859410399506 | 0.01278692581993412 |
| CTA | 0.00711506382014193 | 0.006504753940567456 |
| TTT | 0.017439126770496537 | 0.016691627895167545 |
| GGG | 0.020970495420331817 | 0.02140429437052679 |
| ACT | 0.014857055418331317 | 0.015128140627774787 |
| TCC | 0.010446724072462167 | 0.010281125672483266 |
| TAC | 0.014001127452596426 | 0.013916604717085782 |
| AGA | 0.0347356547296386 | 0.03567341437438628 |
| TCA | 0.01252720261528702 | 0.012377306106860953 |
| GCA | 0.017834699999856814 | 0.01810931634701613 |
| TGC | 0.01907767601664876 | 0.01844420945173924 |
| CTG | 0.01126940836487511 | 0.009318014424929666 |
| ACA | 0.020331959811006276 | 0.02071605856662311 |
| TGG | 0.023797429679653485 | 0.023000057025766765 |
| GGC | 0.021384692427595144 | 0.021731790590543233 |
| GAA | 0.03265659646702997 | 0.0336117112455764 |
| GCT | 0.019650121003653213 | 0.019973284516907094 |
| GGT | 0.02172734736720994 | 0.022157071788892212 |
| CAC | 0.008089816604964814 | 0.00818431077706696 |
| GCC | 0.018490421809010442 | 0.018750251732391605 |
| CTT | 0.0102759693487574 | 0.009061084872546397 |
| TCT | 0.015693984767913186 | 0.014092709709528601 |
| CAG | 0.012951964799886029 | 0.013035104615850003 |
| GGA | 0.03108645622094747 | 0.031677415558334296 |
| CAT | 0.011118987465903474 | 0.01117352137881731 |
| AAA | 0.04150318656882998 | 0.04231364266799029 |
| TAT | 0.02228981595129152 | 0.022247818992847505 |
| TAA | 0.023798416978773242 | 0.023322130156142467 |
| TGA | 0.02834572643664702 | 0.027359521822232467 |
| CAA | 0.011875937590516394 | 0.011820625195415869 |
| TGT | 0.027232638370220368 | 0.026994055138925575 |
| CCC | 0.010055248465641091 | 0.010165488543163232 |
| CCT | 0.01260279650814785 | 0.01272147209025231 |
| CCA | 0.013099524144093254 | 0.013125198379449421 |

### Slide 6
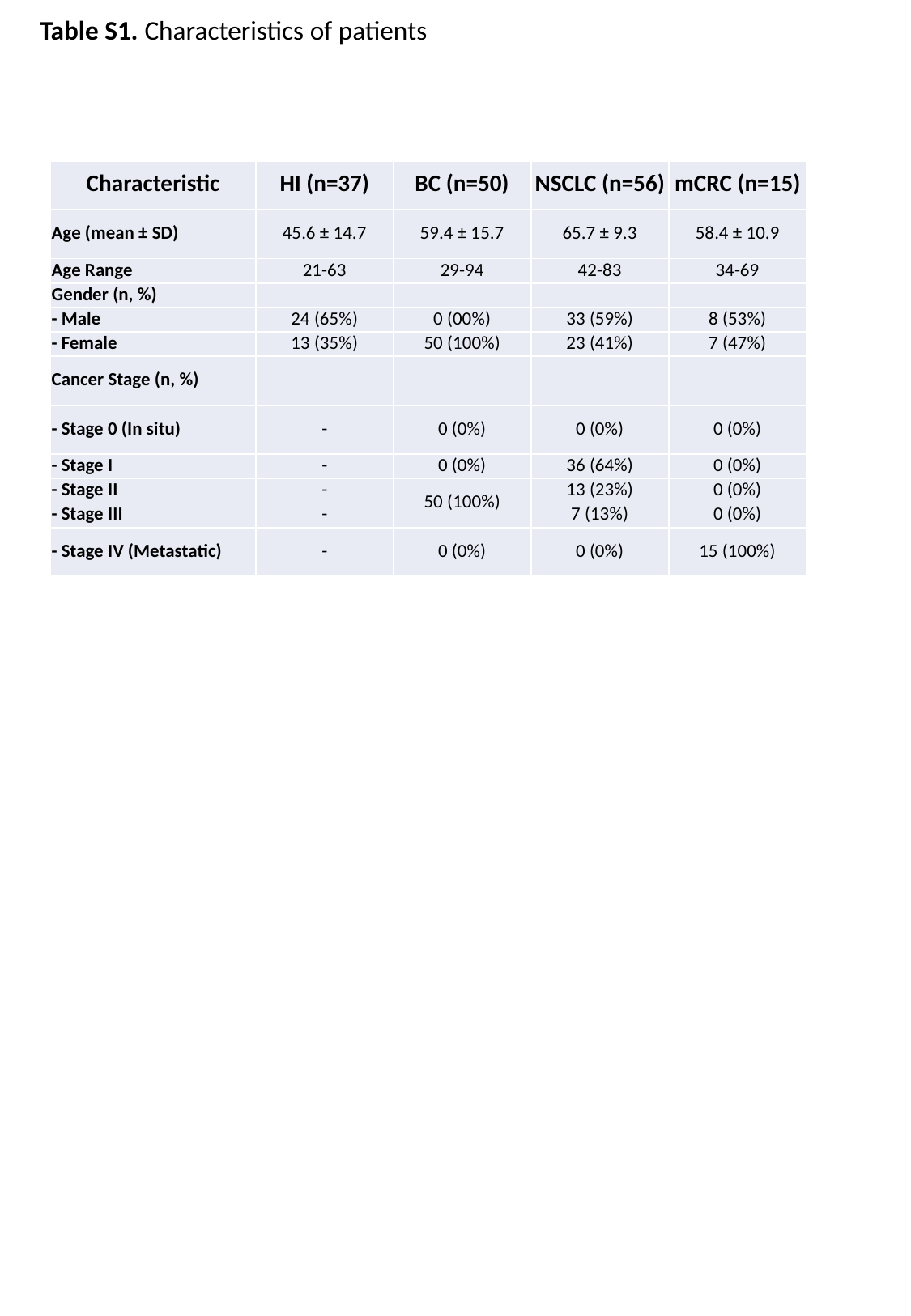

Table S1. Characteristics of patients
| Characteristic | HI (n=37) | BC (n=50) | NSCLC (n=56) | mCRC (n=15) |
| --- | --- | --- | --- | --- |
| Age (mean ± SD) | 45.6 ± 14.7 | 59.4 ± 15.7 | 65.7 ± 9.3 | 58.4 ± 10.9 |
| Age Range | 21-63 | 29-94 | 42-83 | 34-69 |
| Gender (n, %) | | | | |
| - Male | 24 (65%) | 0 (00%) | 33 (59%) | 8 (53%) |
| - Female | 13 (35%) | 50 (100%) | 23 (41%) | 7 (47%) |
| Cancer Stage (n, %) | | | | |
| - Stage 0 (In situ) | - | 0 (0%) | 0 (0%) | 0 (0%) |
| - Stage I | - | 0 (0%) | 36 (64%) | 0 (0%) |
| - Stage II | - | 50 (100%) | 13 (23%) | 0 (0%) |
| - Stage III | - | | 7 (13%) | 0 (0%) |
| - Stage IV (Metastatic) | - | 0 (0%) | 0 (0%) | 15 (100%) |

### Slide 7
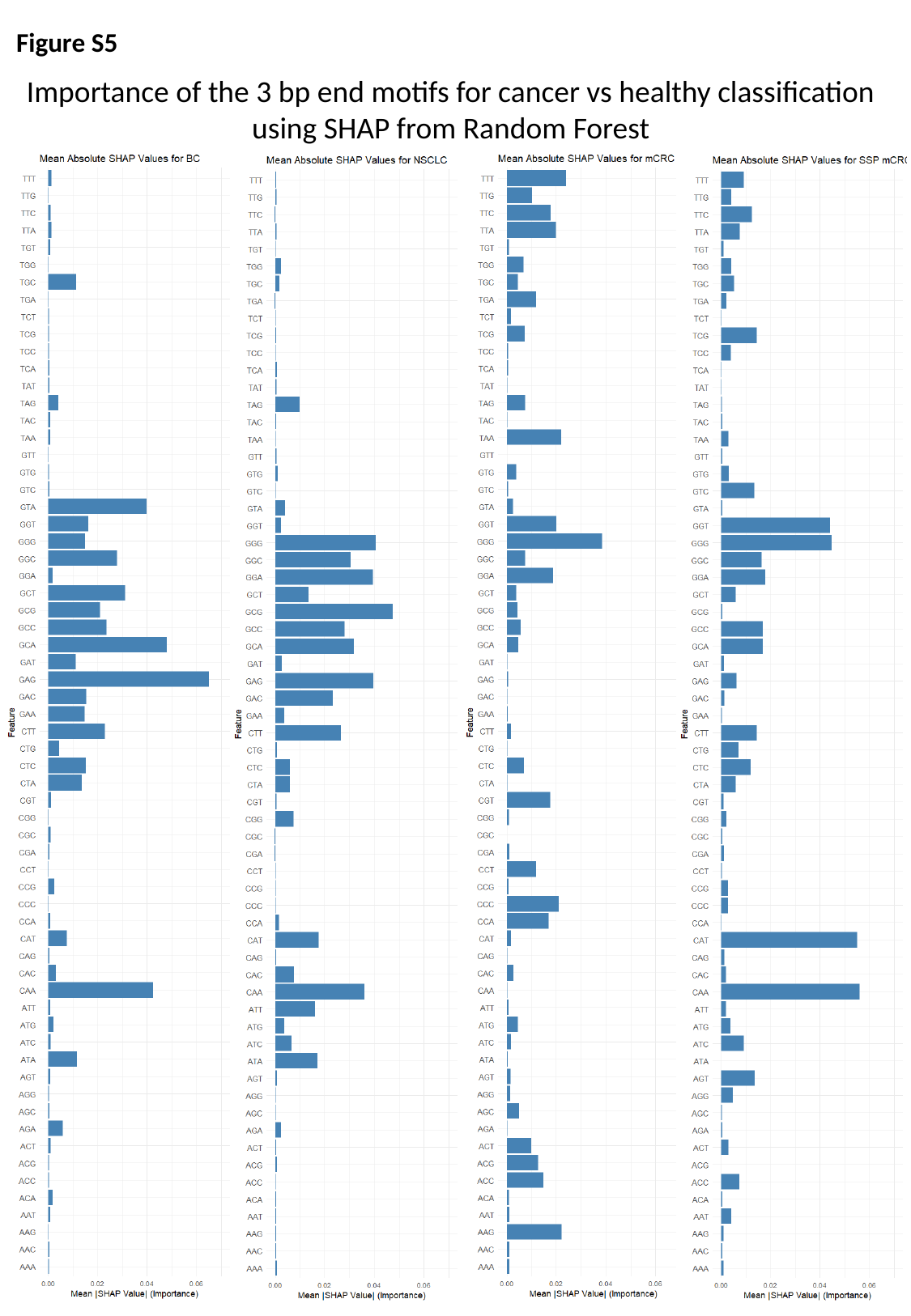

Figure S5
Importance of the 3 bp end motifs for cancer vs healthy classification using SHAP from Random Forest
